# Spatiotemporal Mapping of Tertiary Lymphoid Structure Heterogeneity Shapes Immune Niches and Clinical Outcomes in Intrahepatic Cholangiocarcinoma

**DOI:** 10.64898/2026.01.02.697439

**Authors:** Liaoliao Gao, Jie Mei, Libing Hong, Yuzhi Jin, Jinlin Cheng, Xuqi Sun, Chuan Liu, Bin Li, Xiaomeng Dai, Bo Lin, Yajie Sun, Peng Zhao, Minshan Chen, Rongping Guo, Shan Xin, Jingping Yun, Inmaculada Martínez-Reyes, Wei Wei, Weijia Fang, Xuanwen Bao

## Abstract

Intrahepatic cholangiocarcinoma (iCCA) is a highly lethal malignancy with limited therapeutic options. The spatial architecture and functional diversity of tertiary lymphoid structures (TLSs) in iCCA remain unclear. Here, we present a multimodal spatial atlas of TLSs and identified Intra-tumoral TLSs (iTLSs) as independent prognostic markers. Bulk proteomic profiling of 214 discovery and 155 validation cases identified a four-tier TLS-based TME classification system and supported development of a TLS-predictive random forest classifier. Imaging mass cytometry revealed that iTLS⁺ tumors harbor structured immune architectures, where M1-like tissue-resident macrophages (RTMs), dendritic cells, and CXCL13⁺ CD4⁺ T cells co-localize to form antigen-presenting neighborhoods (apc-CNs) spatially coupled to TLS core regions (TLScore-CNs). Single-cell spatial transcriptomics further resolved 61 TLSs into 14 spatial niches and defined a pseudo-temporal maturation continuum: aggregated, activated, and post-activated. Intra-niche communication, primarily mediated by ifnCAFs, iCAFs, and CXCL12⁺ macrophages, evolved dynamically with maturation. Single-nucleus RNA-sequencing combined with Tangram-based spatial mapping revealed CXCL12⁺ macrophages and iCAFs forming a peripheral band in aggregated TLSs, while ifnCAFs infiltrated TLS interiors during activation. These findings define TLS heterogeneity and provide insights for stroma-directed immunotherapy.

**Teaser:** An atlas of tertiary lymphoid structures reveals immune-stroma interactions in Intrahepatic cholangiocarcinoma.

## Introduction

Intrahepatic cholangiocarcinoma (iCCA), the second most common primary liver malignancy, is marked by aggressive clinical behavior and pronounced molecular heterogeneity, posing significant challenges for effective therapeutic management (*1*). Despite the advent of PD-(L)1 blockade and targeted agents, clinical benefits remain limited to specific subsets of patients (*2*), underscoring the need for a deeper understanding of tumor microenvironmental (TME) features that could inform more precise therapeutic stratification.

Tertiary lymphoid structures (TLSs) have emerged as key immunological components within the TME, with accumulating evidence across multiple cancer types supporting their role in antitumor immunity and their predictive value for immunotherapy response (*3*). In iCCA and related hepatobiliary malignancies, TLSs exhibit striking heterogeneity in both spatial distribution and maturation status, with functional consequences for local immune responses (*4–8*).

Specifically, intratumoral TLSs—particularly follicular structures containing germinal centers—have been associated with improved patient survival and enhanced recruitment of CXCL13⁺ CD4⁺ T cells, reflecting a capacity to support productive T–B cell interactions (*9*). In contrast, peritumoral TLSs may contribute to immunosuppression via Treg enrichment, and immature aggregates (AGG) often lack prognostic significance, highlighting the functional divergence of TLS subtypes.

This prognostic paradigm of TLS heterogeneity is recapitulated in other malignancies. In hepatocellular carcinoma, immature TLSs can be further stratified into conforming and deviating forms, with only the former exhibiting responsiveness to immunotherapy on par with mature TLSs (*10*). Similarly, in high-grade serous ovarian cancer, the spatial maturity of TLSs varies markedly across anatomical sites, with more developed structures correlating with increased immune activity and improved outcomes (*11*). These findings converge on the notion that the spatial architecture, developmental trajectory, and immune functionality of TLSs collectively determine their clinical relevance and therapeutic implications.

Despite these insights, a systematic, high-resolution atlas of TLS heterogeneity in iCCA has yet to be established. To address this, we employed an integrative spatial multi-modal approach, combining bulk and spatial proteomics, single-nucleus RNA sequencing (snRNA-seq), and imaging mass cytometry (IMC) with single-cell spatial transcriptomics. Through this spatial multi-modal framework, we identified four TLS-based TME subtypes—Desertic, Excluded, Immature, and Structured—defined by distinct TLS morphologies and spatial features. Among them, the Structured subtype, characterized by intratumoral TLSs (iTLS), emerged as an independent prognostic marker.

Subtype-specific proteomic signatures enabled construction of a predictive random forest classifier, while paired IMC provided high-resolution maps of spatial architectures. Single-cell spatial transcriptomics further delineated the molecular identity of TLSs, uncovering subtype-specific cellular compositions and pseudo-time dynamic remodeling of spatial niche networks within the TME. snRNA-seq provided active signaling pathways and enabled high-resolution spatial mapping, revealing subtype-specific transcriptional programs. Together, these findings establish a multidimensional framework for understanding spatiotemporal TLS diversity in iCCA, offering insights into its prognostic relevance and potential therapeutic leverage points.

## Results

### Comprehensive Landscape of TLS Distribution and Classification in iCCA

We conducted a spatial and morphological assessment of TLSs in tumor samples from 214 patients with iCCA (Figure 1A). Histological evaluation and staining with CD20, CD3, CD21, BCL6, and Ki67 were used to classify TLSs into three distinct morphologies: aggregates (Agg), primary follicles (FL1), and secondary follicles (FL2) (Figure 1B–C, Figure S1A-B). Spatial profiling showed that 22.2% of TLSs were located intratumorally, with the remainder residing in the peri-tumoral stroma. Density distribution analysis revealed a strong positional preference within 5 mm of the tumor boundary (Figure 1D). Morphological mapping using Sankey diagrams indicated distinct configurations between intra-tumoral (iTLS) and peri-tumoral (pTLS) compartments (Figure 1E). iTLS were detected in 29% patients with mature TLS (FL1 and FL2) exsiting in most cases, while pTLS were observed in 77% patients with immature (Agg) and mature TLS evenly evaluated.

**Figure 1.**
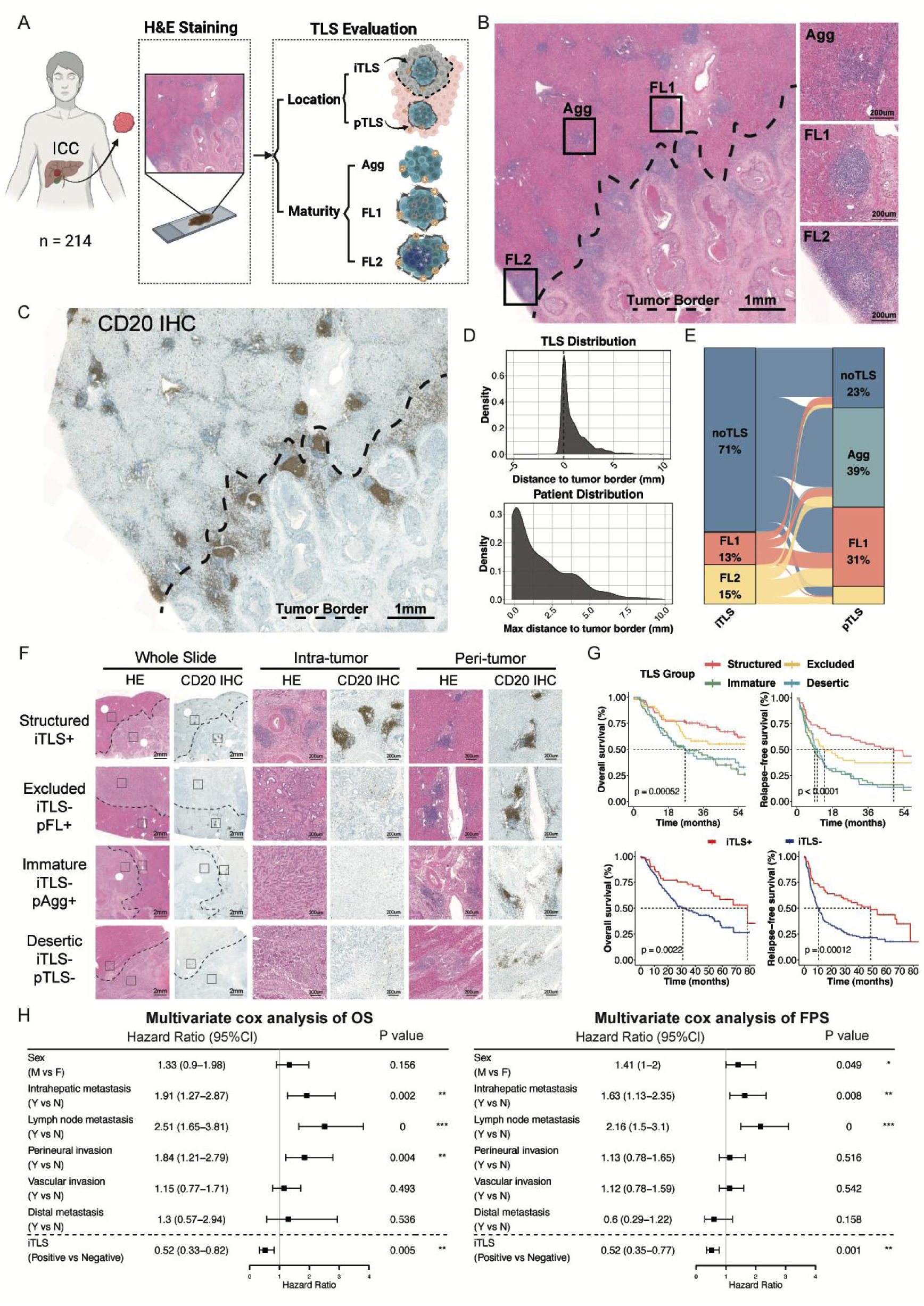
Comprehensive evaluation of TLS distribution, morphology, and prognostic classification in iCCA. (A) Schematic overview of TLS identification, spatial annotation, morphological classification, and prognostic stratification in intrahepatic cholangiocarcinoma (iCCA). (B) Representative hematoxylin and eosin (H&E) stained images showing the three morphological subtypes of tertiary lymphoid structures (TLS): lymphoid aggregates (AGG), primary follicles (FL1), and secondary follicles (FL2). (C) Representative immunohistochemical staining of CD20 highlighting B-cell-rich TLS structures, supporting morphological classification. (D) Density plot showing the spatial distribution of TLS relative to the tumor boundary, demonstrating predominant localization within 5 mm from the tumor edge. (E) Sankey plot illustrating the distribution of TLS morphological subtypes across intra- and peri-tumoral regions. (F) Histological and immunohistochemical images (H&E and CD20 staining) demonstrating the criteria used to define the four-tier TLS classification: Structured (mature intratumoral TLS), Excluded (iTLS– & pFL+), Immature (iTLS– & pAGG+), and Desertic (iTLS– & pTLS–). (G) Kaplan–Meier survival curves showing overall survival (OS) and progression-free survival (PFS) across TME subtypes, indicating significant prognostic stratification. (H) Forest plots from multivariate Cox regression analysis demonstrating iTLS presence as an independent prognostic factor for both OS and PFS.

Based on these patterns, we devised a four-tier TLS-based TME classification system comprising: the Structured subtype (iTLS^+^), Excluded (iTLS^−^/pFL^+^), Immature (iTLS^−^/pAgg^+^), and Desertic (iTLS^−^/pTLS^−^) phenotypes (Figure 1F). This classification showed significant prognostic value, with differences in overall survival (OS, *p* = 0.00052) and progression-free survival (PFS, *p* < 0.0001) across subtypes (Figure 1G). Notably, patients with Structured TLSs exhibited significantly improved outcomes (OS *p* = 0.0037; PFS *p* = 0.00018). In multivariate analysis adjusted for clinical covariates, the presence of iTLS was independently associated with better OS (HR = 0.52, 95% CI: 0.33–0.82, *p* = 0.005) and PFS (HR = 0.52, 95% CI: 0.35–0.77, *p* = 0.001) (Figure 1H).

### Proteomic Profiling Reveals TLS-Associated Immune Subtypes

To investigate the molecular correlates of TLS heterogeneity, we performed multi-omics profiling across a discovery cohort (n = 214) and additional validation cohorts, including spatial proteomics (n = 155), single-nucleus RNA sequencing (snRNA-seq, n = 14), and spatial transcriptomics (n = 6). Mass spectrometry–based bulk proteomics enabled construction of a TLS-predictive random forest classifier, which was subsequently validated using expansion-gel spatial proteomics (n = 155) (Figure 2A). To allow spatially matched validation, expansion-gel spatial proteomics and imaging mass cytometry (IMC) were performed on adjacent tissue sections from the same specimens. Regions of interest (ROIs) were precisely co-registered across modalities to ensure that spatial proteomic data and single-cell spatial data were derived from the same anatomical context. IMC was used to delineate the single-cell spatial architectures of TME subtypes, while 10x Xenium whole-slide spatial transcriptomics enabled comprehensive TLS profiling at tissue scale with subcellular resolution. In parallel, snRNA-seq profiles were applied to spatially mapped onto Xenium spatial transcriptomic coordinates for high-resolution projection and uncover subtype-specific transcriptional programs.

**Figure 2.**
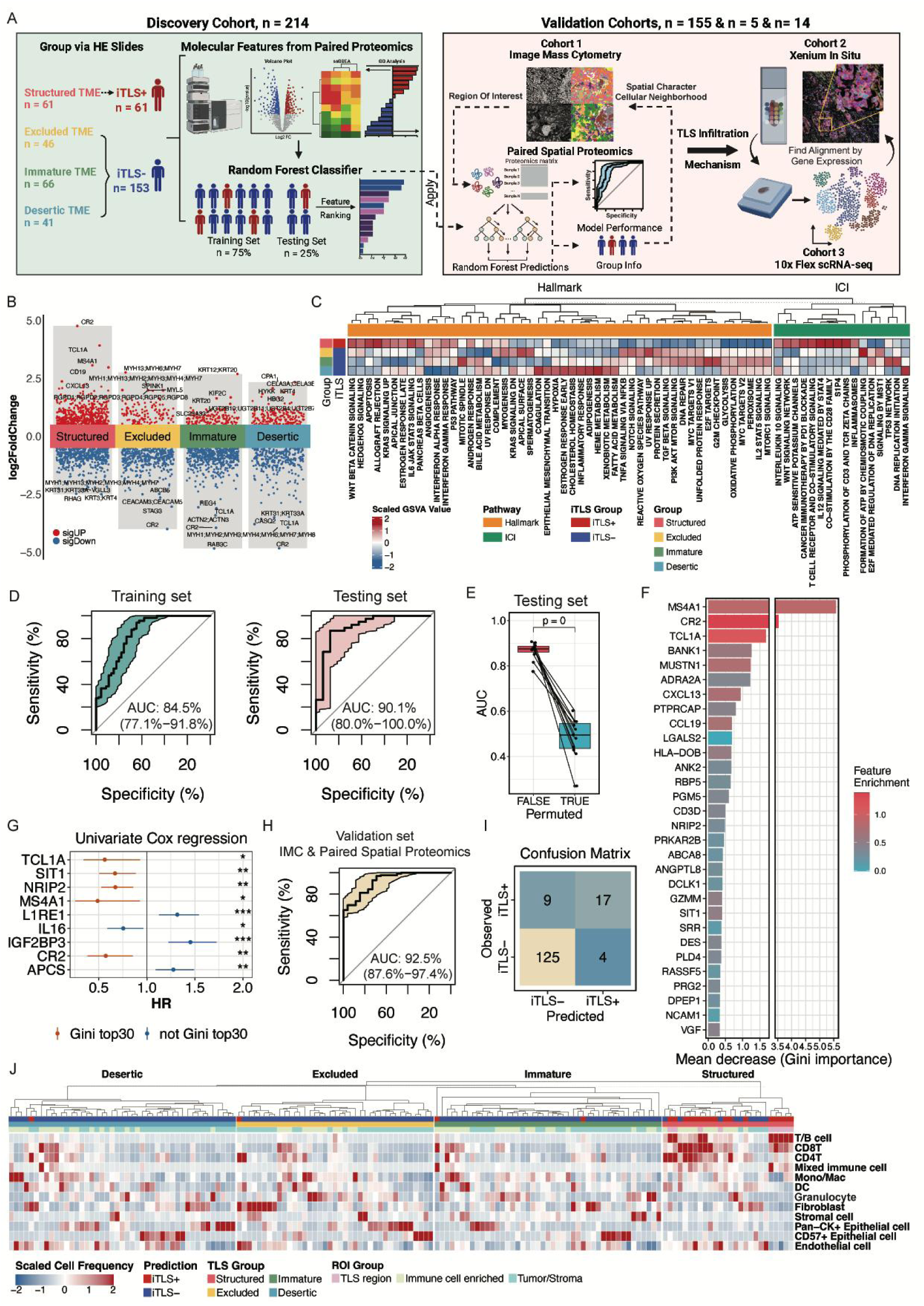
Proteomic and spatial characterization of TLS-associated tumor microenvironment subtypes in iCCA. (A) Overview of the integrated spatial multi-omics strategy used to characterize TLS heterogeneity in iCCA, including bulk proteomics, imaging mass cytometry (IMC), single-nucleus RNA sequencing (snRNA-seq), and spatial transcriptomics (10x Xenium). (B) Volcano plot showing differentially expressed proteins between TME subtypes identified in the discovery cohort (n = 214). Notably, Structured tumors show upregulation of lymphoid markers such as CR2, CD19, MS4A1, and CXCL13, while Desertic tumors upregulate epithelial programs (e.g., KRT4). (C) Heatmap of pathway enrichment scores reveals immunoactive signaling in Structured subtype (e.g., CD28 co-stimulation, S1PR4, IL-12 response) and hypoxia/EMT/coagulation programs in Desertic subtype. (D) Performance of the TME subtype prediction model using a random forest classifier trained on proteomic data, showing training and testing AUCs (84.5% and 90.1%, respectively; p < 0.001). (E) Boxplot comparing AUC values of the random forest model in the testing cohort under true versus permuted TLS labels, demonstrating significant performance drop in permuted conditions and confirming the model’s specificity. (F) Feature importance analysis from the random forest model, highlighting top discriminative proteins including MS4A1, CR2, TCL1A, BANK1, MUSTN1, ADRA2A, and CXCL13. (G) Univariate survival analysis of the top-ranked features and others from TLS-prediction model, showing significant prognostic associations. (H) ROC curve demonstrating external validation performance of the TLS prediction model in an independent iCCA cohort (n = 155) using expansion-gel-based spatial proteomics (AUC = 92.5%). (I) Confusion matrix evaluating prediction accuracy in the validation cohort, confirming high classification performance. (J) Heatmap generated from IMC data showing cellular composition and spatial immune architecture across TME subtypes. Structured tumors feature dense, organized lymphoid aggregates; Excluded and Immature subtypes display compartmentalized immune patterns; Desertic tumors exhibit immune-depleted, stromal-dominant landscapes.

Distinct proteomic signatures were identified across TLS-based TME subtypes (Figure 2B). Structured TLSs were enriched for lymphoid-associated molecules, including CR2, CD19, and MS4A1, along with chemokines such as CXCL13. In contrast, Desertic tumors exhibited increased expression of epithelial and metabolic markers, notably KRT4. Pathway enrichment analysis highlighted subtype-specific immune activity: Structured TLSs showed enhanced immune activation, including CD28 co-stimulation, S1PR4 signaling, and IL-12 response pathways, whereas Desertic tumors exhibited dominant signatures of hypoxia, epithelial–mesenchymal transition (EMT), and coagulation (Figure 2C).

To further validate these subtype-specific immune landscapes, we applied a previously published TLS-related gene signature (*12*) (Supplement table S1) to derive a proteomics-based TLS score. Structured tumors exhibited the highest TLS scores, while Desertic tumors had the lowest (Figure S2A), supporting the subtype classification. GO analysis of differentially expressed proteins among the four subtypes revealed that Structured tumors were enriched for immune-activated pathways—such as B cell receptor signaling, leukocyte proliferation, T cell selection, and immune homeostasis—while Desertic tumors showed upregulation of coagulation, reactive oxygen species metabolism, complement activation, and extracellular matrix remodeling (e.g., cell–substrate adhesion) (Figure S2B). Volcano plot and enrichment analysis highlighted functional immune divergence: iTLS⁺ tumors displayed activation of antigen receptor–mediated signaling, T cell receptor signaling, regulation of B cell proliferation and chemokine signaling (Figure S2C–D). In contrast, iTLS⁻ tumors showed upregulation of ECM reorganization, metabolic reprogramming, and cell cycle–related pathways, including p53 signaling (Figure S2E).

Given the pronounced survival advantage observed in iTLS⁺ tumors, we next asked whether proteomic profiles could effectively distinguish iTLS⁺ from iTLS⁻ samples. We therefore constructed a random forest classifier to predict iTLS status based on protein expression features derived from the discovery cohort. The dataset was randomly split 3:1 into training and testing sets and evaluated using five-fold cross-validation. The model achieved a training AUC of 84.5% and a testing AUC of 90.1% (p < 0.001) (Figure 2D), with permutation control confirming specificity (Figure 2E). Top discriminative proteins included MS4A1, CR2, TCL1A, BANK1, ADRA2A, and CXCL13 (Figure 2F), many of which were also prognostically significant in univariate survival analyses (Figure 2G). External validation in an independent cohort using expansion-gel spatial proteomics combined with IMC achieved an AUC of 92.5%, further supporting model generalizability (Figure 2H–I).

In validation cohort, IMC-based mapping first revealed distinct major cell populations and their finer subdivision into subsets based on specific markers, as shown in Supplementary Figure S3A-C. These analyses identified immune cell subsets and their spatial distribution across the tumor microenvironment. The characterization of these populations allowed us to further investigate the immune landscape in iCCA and correlate it with the TME subtypes. Structured subtypes showed organized lymphoid aggregation, while Excluded and Immature subtypes displayed compartmentalized immune architecture. Desertic tumors exhibited sparse immune infiltration dominated by stromal and epithelial cells (Figure 2J). Notably, the structured subtype revealed a significant clustering of T and B cells within the TLS region ROI, highlighting the organized immune architecture characteristic of mature TLS (Figure 2J and Figure S3D).

### Single-cell Spatial Architecture of TLS-Enriched Tumor Microenvironments

To investigate how immune spatial architecture differs across TLS-defined TME subtypes, we performed IMC to resolve single-cell spatial features within representative tumor regions. IMC was applied to rigorously selected ROIs representing TLSs, immune-infiltrated areas, and stromal compartments (Figure 3A). Survival analysis at whole-slide level in the validation cohort recapitulated subtype associations with OS (*p* = 1e^-04^) and confirmed prognostic value of iTLS presence (*p* = 0.00011) (Figure 3B).

**Figure 3.**
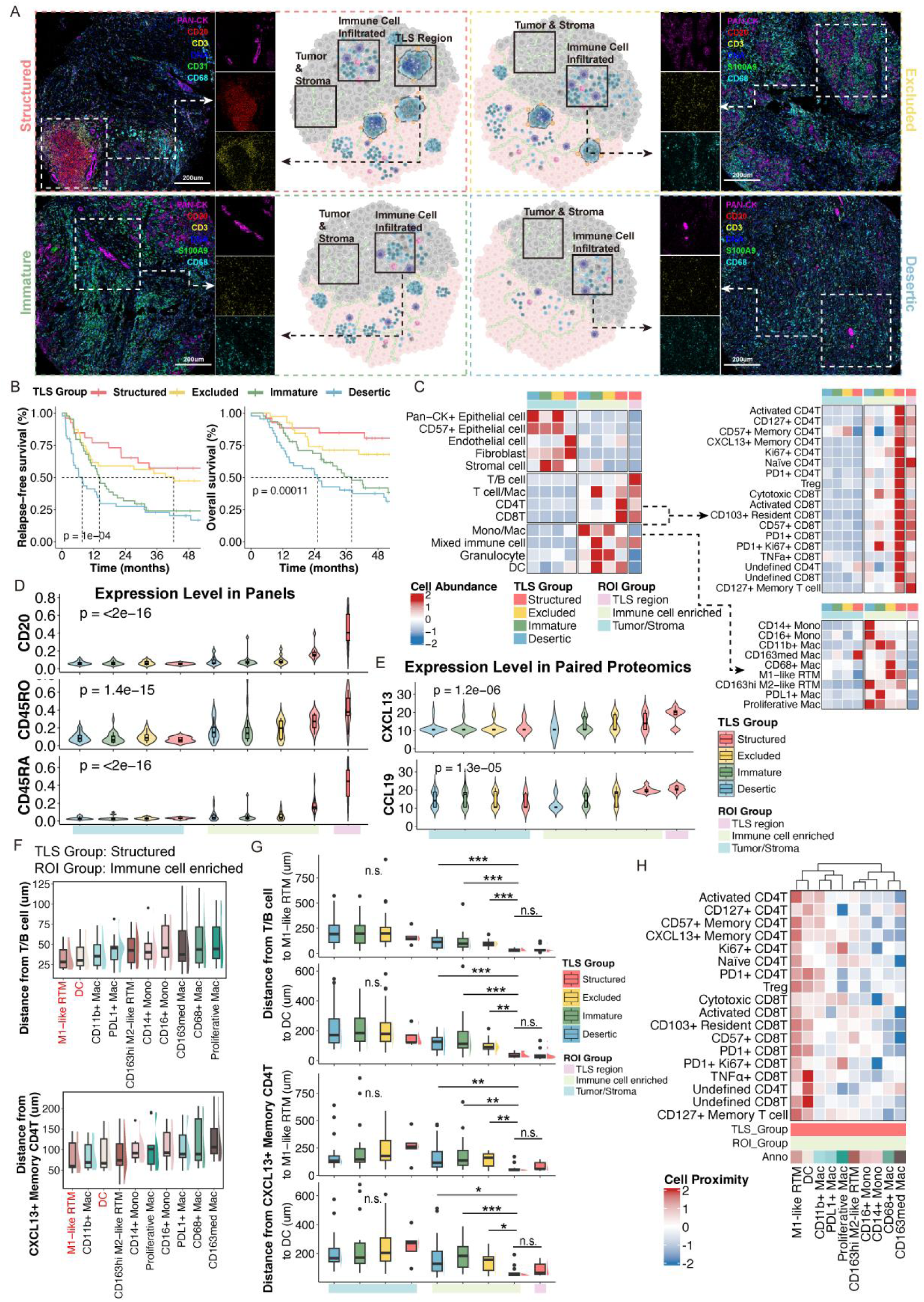
Single-cell spatial architecture of TLS-enriched tumor microenvironments in iCCA. (A) Workflow of imaging mass cytometry (IMC) analysis with region-of-interest (ROI) segmentation in TLS, immune-infiltrated, and tumor/stroma regions across iCCA samples. (B) Kaplan-Meier survival analysis in the validation cohort showing significantly improved overall survival across TME subtypes (p = 1e-04) and between iTLS^+^ and iTLS^−^ groups (p = 0.00011). (C) Heatmap displaying the relative abundance of immune cell subsets across spatial compartments. Structured TLS regions are enriched for CXCL13^+^ TFH-like memory CD4^+^ T cells, naïve CD4^+^ T cells, PD-1^+^ CD4^+^ T cells, and CD20^+^ B cells. (D) Boxplots showing IMC-derived expression levels of key markers (CD20, CD45RO, CD45RA), revealing spatial gradients with highest expression in TLS regions. (E) Violinplot of expansion-gel based spatial proteomics data demonstrating compartment-specific expression of TLS-associated chemokines (CXCL13, CCL19), with enrichment in TLS regions. (F) Boxplots quantifying mean spatial distances between M1-like resident tissue macrophages (RTMs) and lymphoid cells (T/B cells and CXCL13⁺ CD4⁺ T cells), highlighting their close physical proximity in TLS regions. (G) Boxplots quantifying distances from T/B cells and CXCL13⁺ CD4⁺ T cells to RTMs and DCs across spatial compartments. A consistent decrease in distance is observed from tumor/stroma to immune-infiltrated zones and further into TLS regions, reflecting increasing spatial organization of immune cell interactions during TLS maturation. (H) Heatmap of pairwise spatial proximities between myeloid populations and CD4^+^ T cell subsets. M1-like RTMs show preferential interactions with CXCL13^+^ memory CD4^+^ T cells, supporting their role as structural anchors in TLS organization.

Structured TME featured densely intermingled T and B cells forming tightly compacted aggregates resistant to sub-clustering, a hallmark of mature TLS organization. Surrounding immune-rich zones were enriched for specialized T cell subsets, including CXCL13⁺ memory CD4⁺ T cells, CD127⁺ and PD1⁺ CD4⁺ T cells, and naïve CD4⁺ T cells (Figure 3C). Expression gradients of CD20, CD45RO, and CD45RA further validated the spatial stratification (Figure 3D), corroborated by spatial proteomics mapping of CXCL13 and CCL19 (Figure 3E).

Spatial proximity analyses highlighted M1-like RTMs as key architectural hubs, forming tight associations with both T/B clusters and CXCL13⁺ CD4⁺ T cells (Figure 3F). Structured tumors exhibited the closest interactions, while Desertic subtypes were characterized by dispersed and poorly connected immune compartments (Figure 3G). DCs also localized near TLSs, though at slightly increased distances compared to M1-like RTMs (Figure 3F–G). Lastly, M1-like RTMs and DCs showed greatest cell proximity to T cell subpopulations than other myeloid cells (Figure 3H). These findings position M1-like RTMs and DCs as organizational anchors in TLS spatial hierarchies.

### TME Subtype-Specific Spatial Context Networks

To further dissect intercellular interactions, we conducted spatial analyses integrating Effective Score (ES), pairwise cell interaction (PCI), and cellular neighborhood (CN) mapping (Figure 4A). M1-like RTMs and DCs showed peak ES in TLS regions of Structured tumors, suggesting their functional compartmentalization within mature TLSs (Figure 4B). PCI mapping confirmed their privileged proximity to CXCL13⁺ CD4⁺ T cells and T/B clusters (Figure 4C).

**Figure 4.**
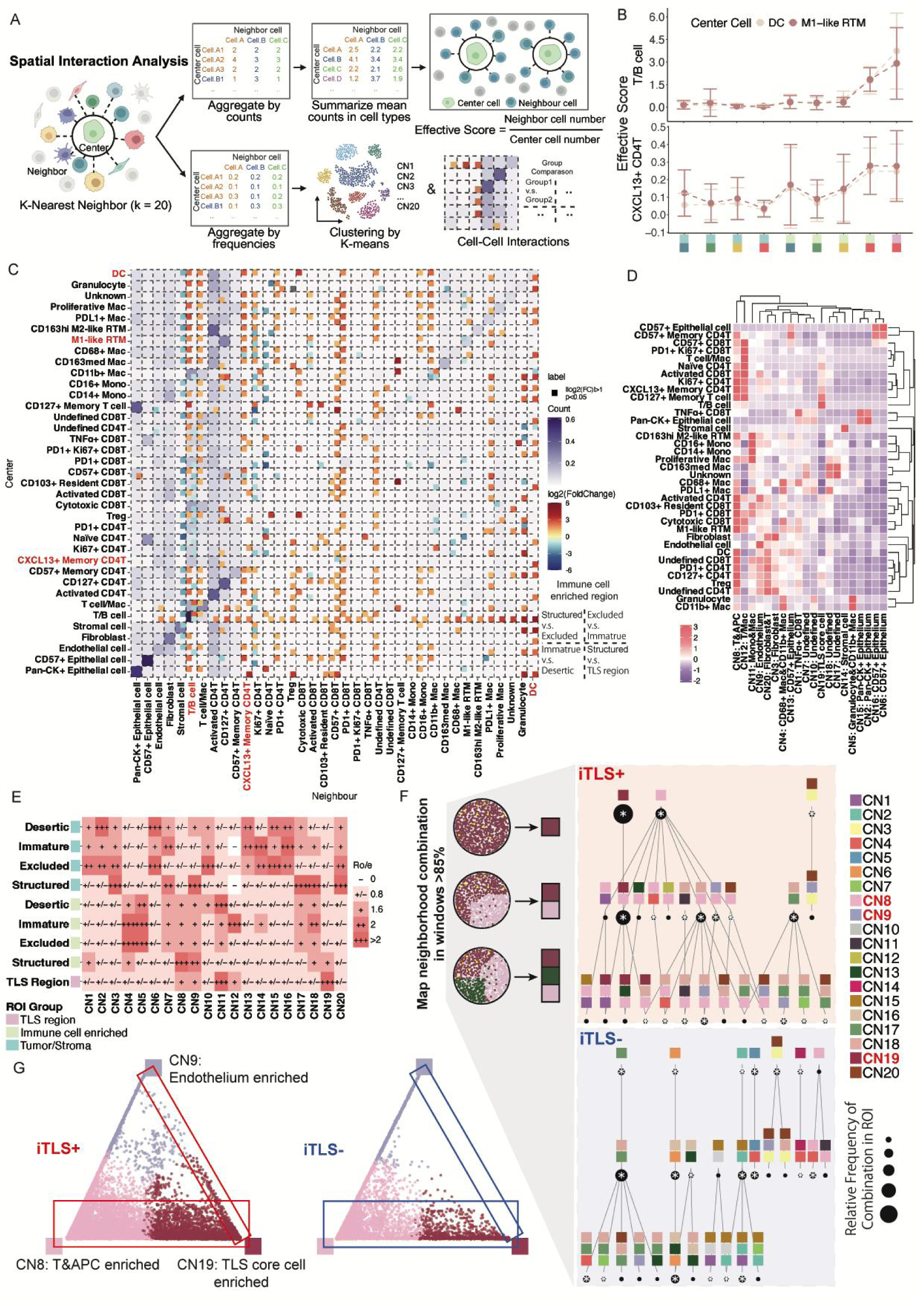
Spatial interaction networks define TLS-associated cellular neighborhoods in iCCA. (A) Schematic workflow for multi-layered spatial analysis integrating Effective Score (ES), pairwise cell interaction (PCI), and cellular neighborhood (CN) profiling. (B) Boxplots showing ES values of M1-like RTMs and DCs across tissue regions, highlighting selective enrichment in TLS regions of structured subtype tumor. (C) Heatmap comparing spatial distances between M1-like RTMs or DCs and CXCL13⁺ CD4⁺ T cells or T/B cell clusters across subtypes. M1-like RTMs and DCs exhibit the shortest interaction distances in structured TLS⁺ tumors. (D) Heatmap depicting relative abundance of major immune cell types across distinct CNs. (E) RO/e analysis showing spatial distribution of CN8 and CN19 across tissue compartments. CN19 is enriched in TLS regions of structured subtype, while CN8 localizes to immune cell-enriched regions of structured subtype. (F) CN interaction analysis illustrating stronger CN8–CN19 interactions in TLS⁺ tumors, particularly in the structured subtype. (G) Triangle plot demonstrating increased co-occurrence of CN8 and CN19 in TLS⁺ tumors, reflecting coordinated spatial coupling within mature TLS regions.

CN analysis revealed two functionally distinct but spatially coupled neighborhoods: CN8, comprising T cells with M1-like RTMs and DCs (apc-CN), and CN19, representing organized T/B cell aggregates (TLScore-CN) (Figure 4D). Regional enrichment analyses demonstrated that CN19 localized specifically to TLS regions, whereas CN8 was predominant in peri-TLS immune cell-enriched zones (Figure 4E). Multicellular neighborhood interfaces may reflect coordinated immune functions within the TME. To quantify and formalize spatial associations between CNs, we applied a spatial context framework adapted from prior studies (*13, 14*). iTLS⁻ tumors exhibited higher degrees of compartmentalization compared to iTLS⁺ tumors (Figure 4F, top row), suggesting a more homogeneous and spatially segregated immune composition. In contrast, CNs such as apc-CN and TLScore-CN showed greater spatial dominance and compartmental stability in iTLS⁺ tumors, indicative of coordinated expansion of APC–T cell–rich TLS regions (Figure 4F, top row). These CNs transformed large tumor regions into structured immune hubs with conserved cell-type compositions.

Spatial windows composed of ≥85% from two CNs were used to define functionally meaningful CN interfaces. iTLS⁺ tumors showed higher frequencies of apc-CN and TLScore-CN co-occurrence compared to iTLS⁻ tumors, where CN pairing appeared more stochastic (Figure 4F, middle row). This suggests that the local compositional balance between these immune neighborhoods may be more functionally relevant than global cell abundance averages.

To further quantify this, we analyzed the relative proportions of apc-CN, TLScore-CN, and other major immune CNs within spatial windows enriched for these interactions (≥85%) (Figure 4G). A dominant interface of apc-CN and TLScore-CN emerged in iTLS⁺ tumors (Figure 4G), which was visibly localized when CNs were spatially mapped (Figure 4G, pink and chestnut red), highlighting their consistent juxtaposition within Structured TLS regions.

These data support a sequential model of TLS organization, wherein antigen presentation and CD4⁺ T cell activation by M1-like RTMs and DCs in CN8 precede the recruitment and spatial organization of lymphocytes into CN19, ultimately establishing mature TLS architecture.

### Single-cell Spatial Transcriptomics Uncovers Molecular States of TLS

Our initial multimodal profiling—combining histopathology, bulk proteomics, and IMC—revealed broad immune heterogeneity in iCCA, with spatial distribution of TLSs correlating with distinct TME landscapes. While IMC identified major TLS-associated neighborhoods, such as dense T/B cell clusters, its limitations in marker depth and spatial throughput precluded fine-resolution dissection of TLS molecular programs. To overcome these constraints, we employed high-content spatial transcriptomics using 10x Genomics Xenium, enabling transcriptome-wide subcellular profiling across intact tissue.

We profiled 433,051 cells from five iCCA samples (4 iTLS⁺, 1 iTLS⁻), with detailed annotation of immune, stromal, and epithelial compartments (Figure 5A–D). TLS-rich zones showed spatially restricted expression of canonical markers (e.g., CXCL13, MS4A1, CR2, CD3E), aligned with cell mask boundaries (Figure 5B). Major cell types and their subtypes—including distinct T cell, B cell, stromal, and myeloid lineages—were resolved at high resolution (Figure 5C–D). Dot plots of marker gene expression validated both major lineages and finer subdivisions (Figure S4A–B).

**Figure 5.**
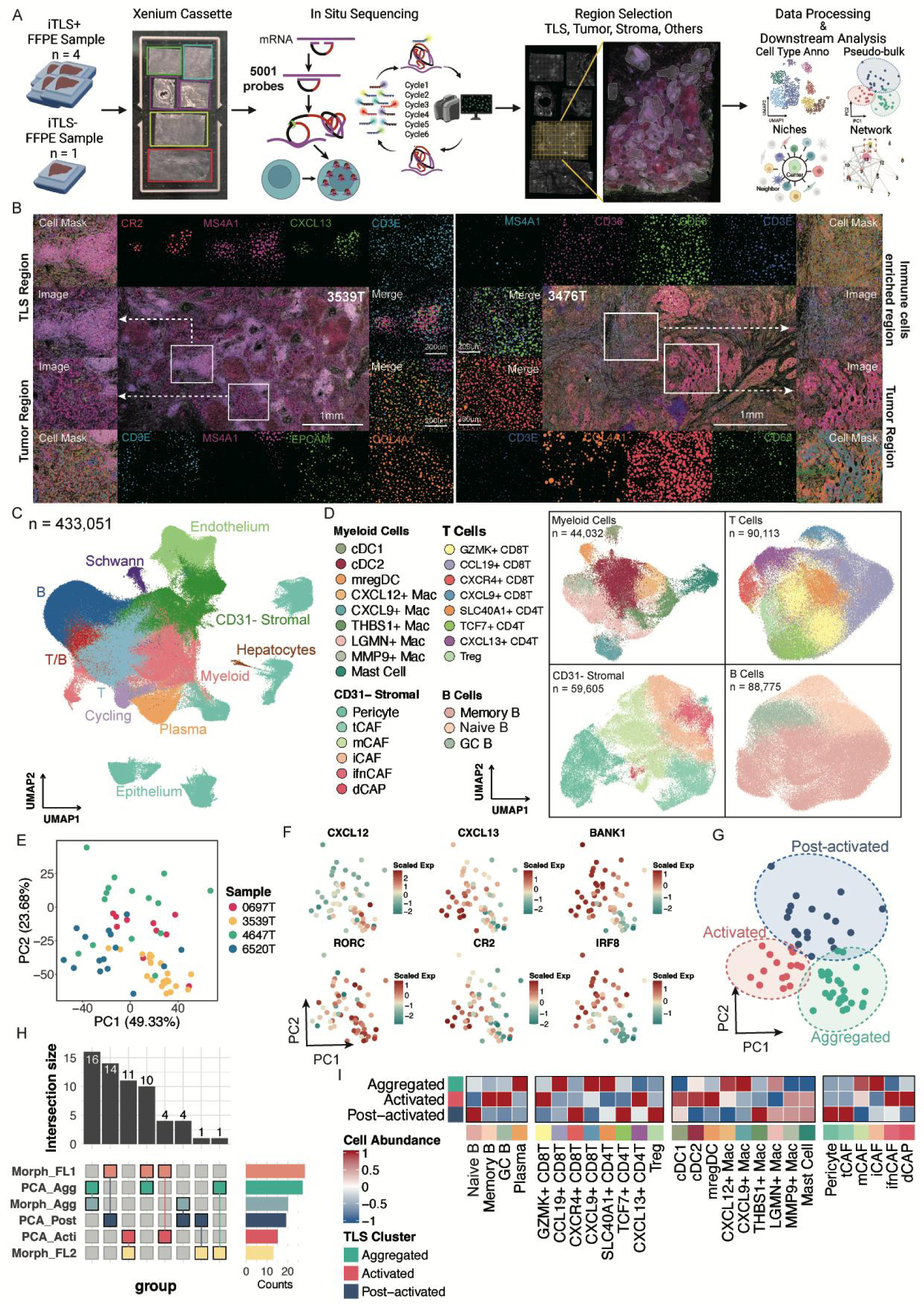
Single-cell spatial transcriptomics reveals TLS molecular heterogeneity and pseudo-temporal subtypes. (A) Schematic overview of the 10x Xenium spatial transcriptomics workflow for profiling TLS at single-cell resolution in iCCA. (B) Representative spatial transcriptomic images from iTLS⁺ and iTLS⁻ samples, showing TLS detection and localization. (C) UMAP embedding of all 433,051 spatially resolved cells colored by major immune and stromal lineages. (D) Sub-UMAPs showing detailed annotation of myeloid cells, T cells, stromal cells, and B cells. (E) Pseudo-bulk PCA treating each of the 61 identified TLS as an individual unit; each dot represents a TLS, colored by patient/sample identity. (F) PCA of TLS colored by expression levels of selected TLS-related markers (e.g., CXCL13, CXCL12, BANK1, RORC, CR2, IRF8), used to define pseudo-temporal states. (G) Pseudo-temporal classification of TLS into three molecular subtypes: Aggregated, Activated, and Post-activated. (H) UpSet plot showing the overlap between molecular subtypes and morphological subtypes (AGG, FL1, FL2), revealing partial concordance but substantial heterogeneity within each morphological class. (I) Heatmap showing differences in cellular composition among TLS molecular subtypes, with Aggregated TLS enriched in plasma cells, Activated TLS enriched in CXCL13⁺ CD4⁺ T cells, and Post-activated TLS enriched in TCF7⁺ CD4⁺ T cells and Tregs.

We next identified 61 TLS structures across samples using combined spatial and transcriptomic criteria. Treating each TLS as a pseudo-bulk unit, we performed PCA on their aggregated gene expression profiles. Principal components PC1 and PC2 captured substantial molecular heterogeneity (Figure 5E), and expression gradients of key TLS-related genes (e.g., CXCL12, CXCL13, BANK1, CR2, RORC, IRF8) revealed continuous, rather than discrete, diversity (Figure 5F; extended in Figure S4C).

Based on PCA positioning and molecular signatures, we stratified TLSs into three pseudo-temporal transcriptional states: Aggregated, Activated, and Post-activated (Figure 5G). While these subtypes partially overlapped with conventional histology (Agg, FL1, FL2), significant intra-class variability highlighted the added granularity of transcriptomic classification (Figure 5H). Abundance analysis of constituent cell types further distinguished these molecular states: Aggregated TLSs were enriched in CXCL12⁺ macrophages, plasma cells, and iCAFs; Activated TLSs featured CXCL13⁺ CD4⁺ T cells; and Post-activated TLSs were characterized by TCF7⁺ CD4⁺ T cells and Tregs (Figure 5I). These findings suggest a coordinated progression from stromal priming to immune activation and eventual regulatory consolidation.

### Spatiotemporal Niche Rewiring and Intercellular Signaling Define TLS Maturation

To elucidate how TLSs mature within the iCCA tumor microenvironment, we systematically mapped 14 spatially resolved cellular niches across the pseudo-temporal trajectory of TLS progression—Aggregated, Activated, and Post-activated. Each TLS stage exhibited distinct niche compositions and organizational architectures (Figure 6A–B). Aggregated TLSs were enriched for niches containing iCAFs and CCL19⁺ CD8⁺ T cells, as well as antigen-presenting niches harboring cDC2 and CXCL12⁺ macrophages. In contrast, Activated TLSs displayed marked expansion of niches enriched in memory B cells, CXCL13⁺ CD4⁺ T cells, IFN-producing CAFs, mregDCs, and GZMK⁺ CD8⁺ T cells, consistent with a transition toward coordinated immune activation. Post-activated TLSs were dominated by regulatory niches composed of TCF7⁺ CD4⁺ T cells, Tregs, and germinal center–like regions populated by naïve and proliferating B cells (Figure S5A-B).

**Figure 6.**
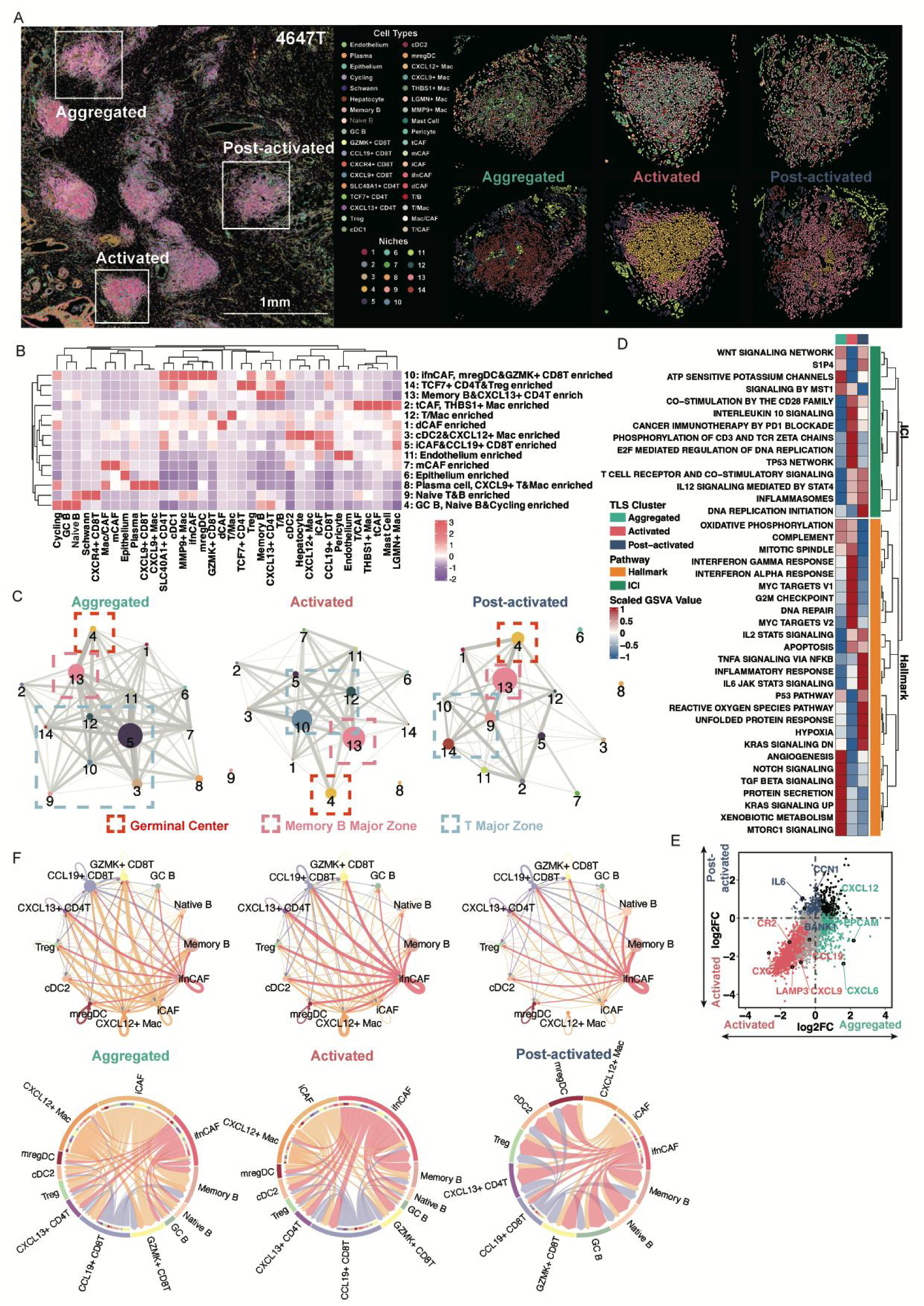
Pseudo-temporal spatial niche remodeling and intercellular signaling across TLS maturation states in iCCA. (A) Schematic of spatial niche analysis across three TLS molecular states (Aggregated, Activated, Post-activated) based on single-cell spatial transcriptomic data. (B) Heatmap displaying the cellular composition of 14 spatial niches, with each niche characterized by specific enrichment of stromal, myeloid, and lymphoid cell subsets. (C) Niche interaction networks reconstructed separately for Aggregated, Activated, and Post-activated TLS, revealing stage-specific topologies. Aggregated TLS show sparse stromal-lymphoid connectivity; Activated TLS exhibit expanded immune coordination; Post-activated TLS display dense regulatory niche interactions. (D) Heatmap of single-sample GSEA (ssGSEA) scores demonstrating hallmark and immune checkpoint-related pathway activities across TLS states. Activated TLS show enrichment for IFN-γ and antigen presentation pathways, while Post-activated TLS are enriched for Treg and immune regulatory signatures. (E) Volcano plot showing differentially expressed genes across TLS molecular states, highlighting key immune regulators and TLS-related chemokines (e.g., CXCL12, IL6, LAMP3, CR2). (F) Intercellular communication networks and CXCL12 signaling dynamics during TLS maturation. Top panels: Niche-specific communication networks in Aggregated (left), Activated (middle), and Post-activated (right) TLS subtypes, showing dominant cell-cell interactions (edges) and key signaling cell populations (nodes). Bottom panels: Corresponding CXCL12 signaling pathways across maturation stages, illustrating dynamic changes in signal-sending cell types (colored nodes) and interaction strengths (edge weights).

To further dissect the spatial coordination of immune compartments, we performed niche-level interaction network analysis, revealing that germinal center and memory B cell zones formed distinct niche–niche interaction hubs at each TLS stage (Figure 6C). In the aggregated subtype, niche connectivity analysis identified significant spatial interactions between iCAF & CCL19⁺ CD8⁺ T cell-enriched niches and both memory B cells & CXCL13⁺ CD4⁺ T cell-enriched niches and cDC2 & CXCL12⁺ macrophage-enriched niches. While the activated subtype exhibited reorganization of the interaction landscape, where memory B cells & CXCL13⁺ CD4⁺ T cell-enriched niches occupied central network positions and demonstrated enhanced connectivity with diverse partners including IFN-producing CAFs, mregDCs, and GZMK⁺ CD8⁺ T cell-enriched niches and other niches, collectively highlighting subtype-specific topological shifts during TLS functional progression. Supplementary Figure S5C further validates this shift by mapping cell–cell network interactions across TME subtypes, complementing niche-level architecture described in Figure 6C.

Hallmark pathway enrichment further demonstrated functional divergence across TLS states (Figure 6D). Aggregated TLSs were enriched for angiogenesis and TGF-β signaling pathways, while Activated TLSs showed prominent interferon responses (IFN-α, IFN-γ), IL-10 signaling, PD-1 pathway activation, and CD3/TCR phosphorylation. Post-activated TLSs exhibited heightened signatures of oxidative stress (ROS pathway), TNF-α signaling via NF-κB, inflammatory responses, and IL6–JAK–STAT3 activation, suggesting immunoregulatory consolidation. Differential expression analysis supported these patterns, with Aggregated TLSs expressing higher levels of CXCL6, Activated TLSs upregulating CR2, CXCL9, and LAMP3, and Post-activated TLSs marked by increased IL6 and CCN1 (Figure 6E).

To investigate intra-niche signaling, we mapped cellular communication networks across TLS maturation states (Figure 6F). We identified strong intra-niche communication circuits where major sender cell types—such as IfnCAF, iCAF, CXCL12⁺ macrophages, and CCL19⁺ CD8^+^ T cells—engaged with recipient cells localized within the same niche microenvironment (Figure 6F, top panel). Furthermore, dynamic shifts were found in dominant signaling cell populations: Aggregated TLSs were characterized by prominent signaling from CXCL12⁺ macrophages and iCAFs, while Activated TLSs showed a transition to IFN-producing CAFs as the primary signal senders (Figure 6F, top panel). Notably, CXCL12-mediated signaling exhibited stage-specific patterns - in Aggregated TLSs, CXCL12 signals originated predominantly from macrophages and iCAFs; during the Activated phase, IFN-producing CAFs became the main CXCL12 source; and in Post-activated TLSs, CXCL12 signaling became more evenly distributed across multiple cell types (Figure 6F, bottom panel). These tightly compartmentalized signaling patterns delineate a spatially organized mechanism whereby niche-specific immune communication orchestrates both microenvironment construction and phase transition during TLS development.

Collectively, these findings define a spatiotemporal trajectory of TLS maturation, transitioning from stromal priming to immune activation and ultimately regulatory consolidation, orchestrated through dynamic niche rewiring and stage-specific intercellular communication networks.

### snRNA-seq and Spatial Mapping Reveal TLS-Associated Immune Topologies

To further validate the spatial and functional signatures identified via IMC and Xenium, we performed single-nucleus RNA sequencing using the 10x Genomics Chromium Flex platform, which offers significantly higher gene detection sensitivity per cell compared to Xenium, across 14 iCCA samples (8 iTLS⁺, 6 iTLS⁻) (Figure 7B). A total of 105,226 cells were profiled, encompassing major compartments including T cells, B cells, myeloid cells, stromal cells, endothelial cells, epithelial cells, cycling cells, and hepatocytes. Refined clustering identified six T cell subsets (e.g., TCF7⁺ CD8⁺ T cells, CXCL13⁺ CD4⁺ T cells), three B cell subsets (naive B, memory B, and germinal center B cells), four stromal subsets (iCAF, ifnCAF, myoCAF, and pericyte), and twelve transcriptionally distinct myeloid populations (e.g., cDC1, cDC2, CD16^+^ monocytes, CXCL12⁺ macrophages) (Figure 7C). Cross-platform validation via dot plots in Supplemental Figure S6A–E confirmed concordance between Xenium and 10x Flex in identifying these subsets, with the latter exhibiting superior gene detection sensitivity (Figure S6A-E).

**Figure 7.**
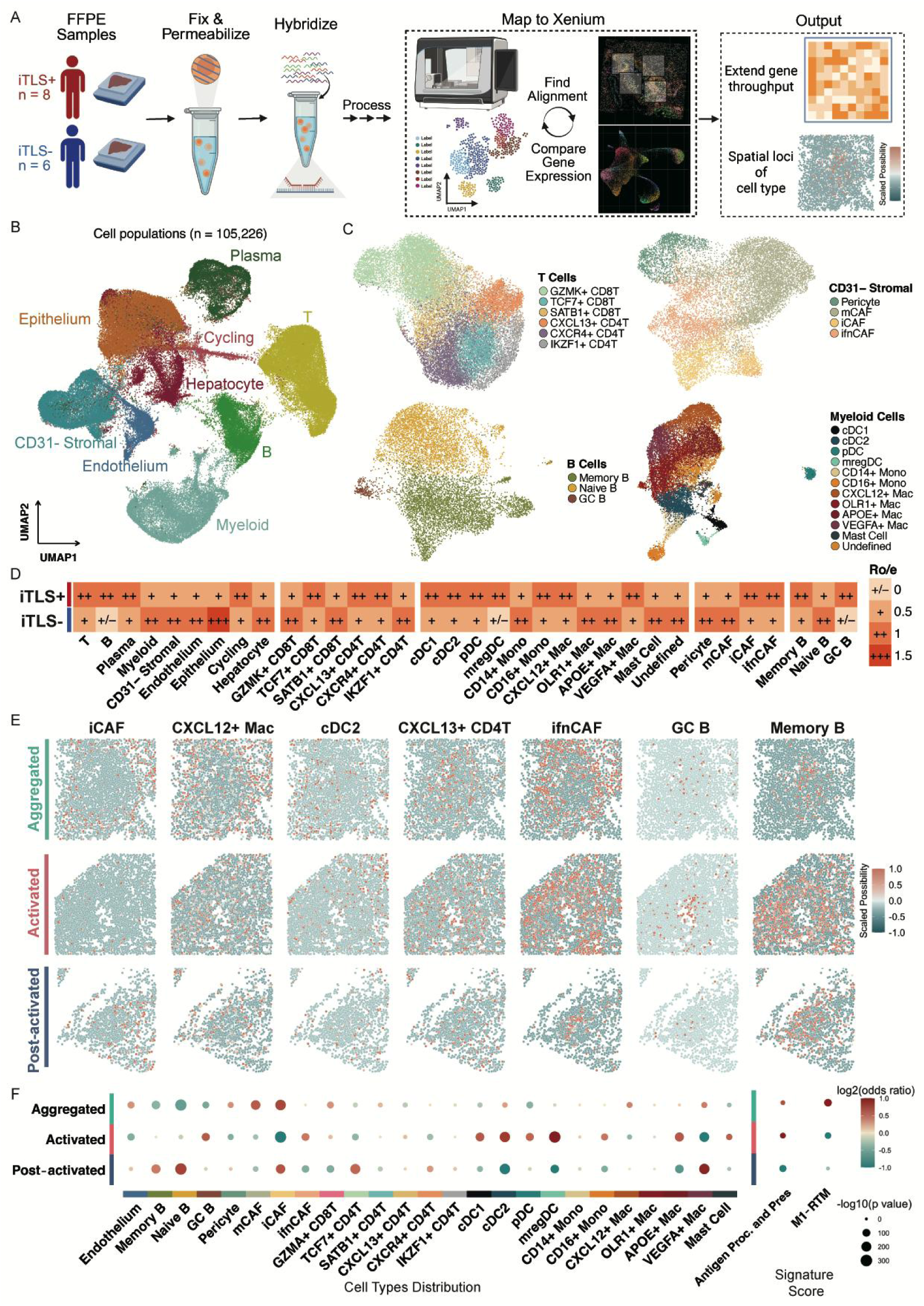
Single-nucleus RNA sequencing and cross-platform spatial mapping reveal TLS-associated immune programs and spatial configurations. (A) Schematic of the integrative workflow combining 10x Chromium Flex single-nucleus RNA sequencing and Tangram-based spatial mapping to Xenium spatial transcriptomics coordinates. (B) UMAP projection of 105,226 nuclei across 14 iCCA samples (8 iTLS⁺, 6 iTLS⁻), showing major cell compartments including T cells, B cells, myeloid cells, stromal cells, endothelial cells, epithelial cells, cycling cells, and hepatocytes. (C) Refined clustering of T, B, myeloid, and stromal populations into functionally distinct subsets, including TCF7⁺ CD8⁺ T cells, CXCL13⁺ CD4⁺ T cells, germinal center B cells, iCAFs, IFNCAFs, and CXCL12⁺ macrophages. (D) Ro/e analysis comparing cellular composition between iTLS⁺ and iTLS⁻ tumors at both major lineage and subset levels. iTLS⁺ tumors are enriched for memory-type and antigen-presenting populations, while iTLS⁻ tumors display elevated epithelial, stromal, and endothelial content. (E) Tangram-projected spatial mapping of Flex-identified subsets onto Xenium spatial transcriptomics coordinates reveals structured localization patterns. CXCL12⁺ macrophages and iCAFs localize to TLS periphery in Aggregated states, while CXCL13⁺ CD4⁺ T cells and memory B cells concentrate in TLS centers in Activated states. (F) Odds ratio (OR) analysis quantifying spatial distribution of mapped cell types across TLS molecular subtypes (Aggregated, Activated, Post-activated), validating subtype-specific enrichment patterns observed in spatial transcriptomic data.

Ro/e analysis revealed preferential accumulation of T cells, B cells, and plasma cells in iTLS⁺ tumors, whereas stromal, endothelial, epithelial, and myeloid cells were more abundant in iTLS⁻ tumors (Figure 7D). At the subset level, immune populations associated with memory and helper functions, including TCF7⁺ CD8⁺ T cells, memory B cells, and CXCL13⁺ CD4⁺ T cells with TFH-like properties, were enriched in iTLS⁺ tumors. Antigen-presenting cells such as cDC1, cDC2, and CXCL12⁺ macrophages were also more prevalent in the iTLS⁺ group. In line with spatial transcriptomics results, stromal subsets including iCAF and ifnCAF showed increased abundance in iTLS⁺ tumors (Figure 7D).

We further dissected the functional polarization of CXCL12⁺ macrophages, which exhibited elevated activities of antigen presentation pathway and M1-related pathway (Figure S6F). Comparative analysis of M1/M2 gene sets confirmed their M1-like phenotype, distinct from other monocyte-derived macrophages (Figure S6G). Notably, these cells scored highest for an M1-RTM signature derived from prior work (Figure S6H), underscoring their role in early TLS initiation.

We next applied Tangram to spatially map the single-nucleus RNA-seq profiles onto Xenium spatial transcriptomic coordinates, thus enabling high-resolution projection of transcriptionally defined cell types and at the same time expanding the gene throughput in spatial tissue architecture (Figure 7E). This integration compensated for Xenium’s limited gene detection by precisely localizing 10x Flex-defined states and their associated pathways. To evaluate the accuracy of spatial mapping between single-nucleus RNA-seq and Xenium spatial transcriptomics, we performed quality control based on Tangram projection metrics. The distribution of training scores indicated overall robust alignment between modalities, with most cells exhibiting high mapping confidence (Figure S7A, left). Sparsity analyses further validated mapping consistency, showing expected relationships between cellular sparsity and spatial signal distribution (Figure S7A, middle and right). Spatial prediction scores were inversely correlated with pixel-level sparsity, yielding a global AUC of 0.704 (Figure S7B), supporting the reliability of Tangram integration. Finally, mapped cell-type compositions across TME subtypes recapitulated biologically coherent patterns demonstrating the spatial fidelity of projection (Figure S7C).

We revealed that iCAF, and CXCL12⁺ macrophages were predominantly enriched in Aggregated TLSs and localized to the peripheral regions of TLS structures, whereas CXCL13⁺ CD4⁺ T cells, memory B cells, and ifnCAFs were more abundant in Activated TLSs. Notably, iCAF and ifnCAF populations formed a ring-like peripheral organization in Aggregated TLSs, which progressively infiltrated into the TLS core in Activated TLSs, reflecting a potential stromal remodeling process.

To further validate the spatial allocation of key cell types across TLS maturation states, we assigned each Xenium-detected cell to its most probable snRNA-seq–defined identity and performed a OR analysis across the 61 TLSs (Figure 7F). The distributions confirmed the expected localization patterns observed in earlier analyses—for example, iCAF, CXCL12⁺ macrophages predominated in Aggregated TLSs, while mregDCs were enriched in Activated TLSs—thereby supporting the spatial validity of the Xenium-based classification framework. The signature scores across mapped cells were further calculated. The M1-RTM signature, derived from macrophages with M1-like tissue-residency properties (*4*), was predominantly enriched in Aggregated TLSs, indicating the early involvement of CXCL12⁺ macrophages in TLS initiation (Figure 7G). In contrast, the APC signature peaked in Activated TLSs, suggesting that robust antigen presentation programs emerge later as TLSs transition into functionally active immune niches.

Together, these integrated results from Xenium and 10x Flex reveal the dynamic spatial reorganization of stromal and immune compartments across TLS developmental states. In combination with IMC-derived spatial CNs, these findings reinforce the model in which antigen presentation and CD4⁺ T cell activation by M1-like RTMs and DCs in early-stage TLS regions precede lymphocyte aggregation and organization into TLScore-CNs, ultimately establishing mature TLS structures. Spatial localization of DCs and CXCL12⁺ macrophages at TLS peripheries in Aggregated states further implicates them as early orchestrators of TLS nucleation.

### Key TLS-Associated Niches Predict Immunotherapy Response

To explore the potential relevance of TLS-associated spatial niches in predicting immunotherapy efficacy, we investigated the prognostic value of five key niches—N3 (cDC2 & CXCL12^+^ Mac enriched), N4 (GC B & Naïve B enriched), N5 (iCAF & CCL9^+^ T & Mac enriched), N10 (ifnCAF, mregDC & GZMK^+^ CD8^+^ T enriched), and N13 (Memory B & CXCL13^+^ CD4^+^ T enriched)—previously implicated in TLS molecular remodeling across the Aggregated to Post-activated continuum. Using three publicly available bulk RNA-seq datasets from immune checkpoint blockade–treated cohorts (GSE91061, GSE126044, and PRJEB23709), which include clinical outcome data from melanoma and non–small cell lung cancer patients, we computed niche-specific signature scores to each cohort. Across all datasets, we observed consistent enrichment of these niche signatures in responders compared to non-responders, indicating their association with favorable therapeutic responses (Figure 8A). Moreover, elevated expression of niche-specific signatures correlated with prolonged overall survival, suggesting that transcriptional features reflective of these spatial niches may serve as conserved biomarkers of effective anti–PD-1–mediated immune engagement (Figure 8B-D). These results suggest that TLS-centric spatial programs may contribute to effective antitumor immunity and serve as predictive markers of immunotherapy benefit.

**Figure 8.**
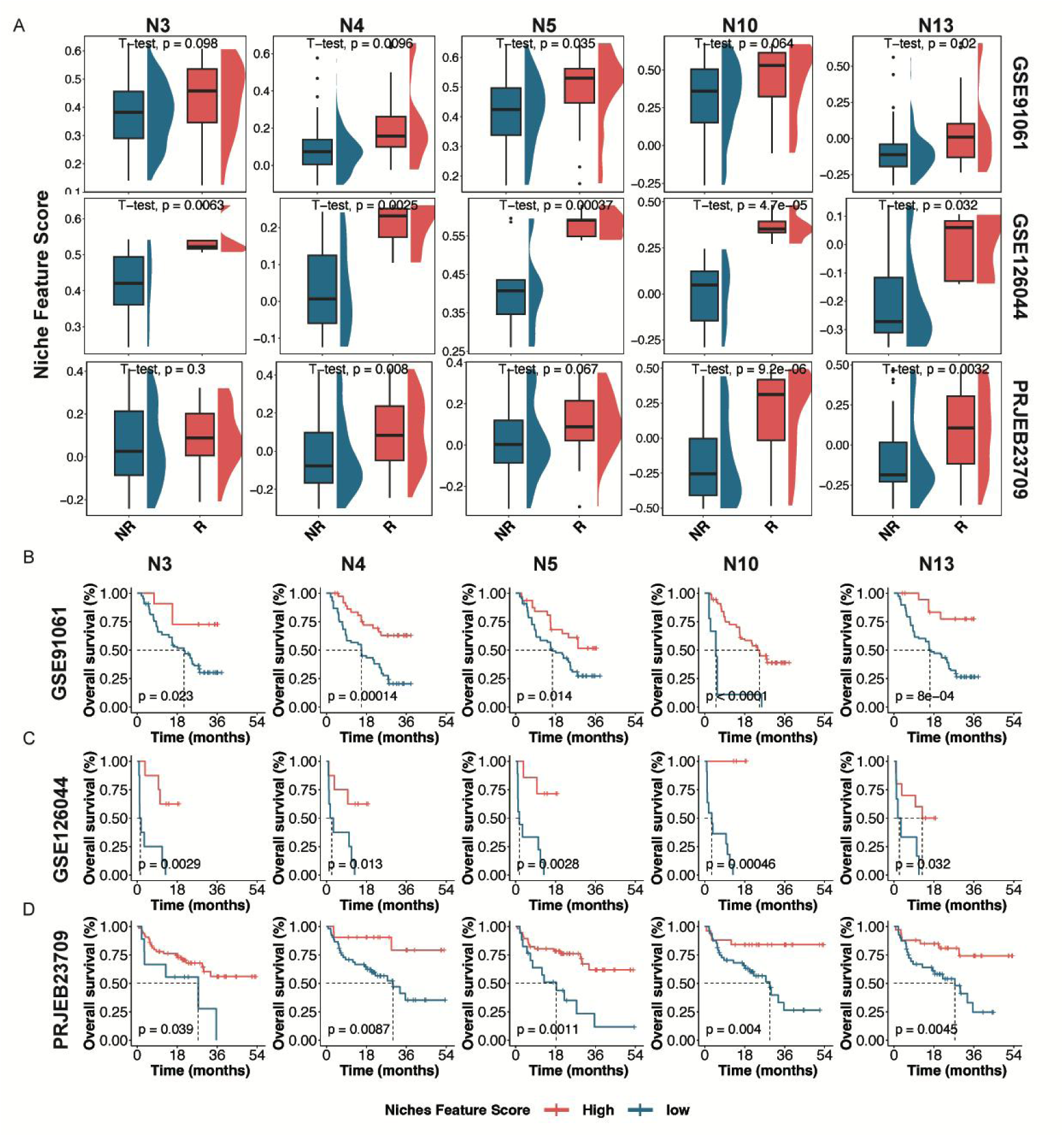
TLS-Associated niches predict immunotherapy benefit and survival outcome in immunotherapy-treated cohorts. (A) Boxplots showing the signature scores of five TLS-associated spatial niches: N3 (cDC2 & CXCL12^+^ Mac enriched), N4 (GC B & Naïve B enriched), N5 (iCAF & CCL9^+^ T & Mac enriched), N10 (ifnCAF, mregDC & GZMK^+^ CD8^+^ T enriched), and N13 (Memory B & CXCL13^+^ CD4^+^ T enriched) in responder versus non-responder groups across three independent immune checkpoint blockade cohorts (GSE91061, GSE126044, PRJEB23709). Signature scores were calculated using GSVA. (B-D) Kaplan–Meier survival curves showing overall survival stratified by the signature score levels of TLS niche 3, 4, 5, 10, and 13 signatures in three independent immunotherapy-treated bulk RNA-seq datasets: (B) GSE91061 (anti–PD-1/anti-CTLA4–treated melanoma and non-small cell lung cancer), (C) GSE126044 (anti–PD-1–treated non–small cell lung cancer), and (D) PRJEB23709 (anti–PD-1/anti-CTLA4–treated melanoma and non-small cell lung cancer). Signatures were computed using GSVA scores of niche-specific genes. p-values were calculated using the log-rank test.

## Discussion

Our study presents the first comprehensive spatial atlas of TLSs in iCCA, unveiling their remarkable molecular and architectural heterogeneity. By classifying iCCA into four distinct immune subtypes based on TLS spatial and morphological features, we propose a novel framework for interpreting tumor–immune interactions. Notably, the identification of the Structured subtype, characterized by the presence of iTLS, as an independent prognostic factor highlights that it is the spatial configuration—rather than the mere presence—of TLSs that dictates clinical outcome. This observation is consistent with findings in other tumor types (*8*), while also revealing iCCA-specific patterns in TLS biology.

To achieve a multidimensional understanding of TLS heterogeneity, we implemented an integrative, multimodal strategy combining bulk and spatial omics platforms. Bulk proteomics enabled large-scale molecular stratification and served as the basis for TME subtype classification via a random forest model. IMC offered single-cell spatial resolution but was limited in multiplexing and tissue throughput. Expansion-based spatial proteomics provided broader cohort-level coverage, albeit at pseudo-bulk resolution. To resolve intra-TLS cellular architecture, we applied single-cell spatial transcriptomics, allowing high-resolution classification of immune and stromal subsets. Given its lower gene detection sensitivity, we further incorporated single-nucleus RNA sequencing to enhance transcriptomic depth and resolve functional programs. Through coordinated use of these complementary technologies, we captured TLS-associated features at multiple molecular and spatial scales—enabling cross-validation of cell types, spatial niches, activation states, and communication networks. This multimodal integration not only reinforces the robustness of our findings but also establishes a scalable framework for decoding complex immune ecosystems in solid tumors.

Our spatial proteomic and transcriptomic analyses underscore the requirement for precise cellular architecture in the formation of functional TLS niches. Specifically, M1-like RTMs, DCs, and T cells form highly organized apc-CN, which spatially interact with TLScore-CN. This architectural coordination recapitulates the microanatomy of lymph nodes, suggesting a conserved paradigm of immune cell orchestration across secondary and ectopic lymphoid sites. The identification of apc-CN as key regulatory hubs within TLS ecosystems offers several mechanistic insights. First, transcriptional analysis via snRNA-seq revealed enrichment of antigen presentation pathways in CXCL12⁺ macrophages (M1-like RTM phenotype) and DCs, implicating their role in local T cell priming. Second, their close spatial proximity to CXCL13⁺ CD4⁺ T cells suggests a function in maintaining TLS-associated chemokine gradients (e.g., CXCL12, CXCL13) (*15, 16*). Third, their interaction networks with TLScore-CN imply potential support for humoral immunity (*17, 18*). These findings complement recent work showing myeloid-dependent TLS activities (*3*), while revealing iCCA-specific organizational patterns.

Leveraging Xenium spatial transcriptomics, we further resolved TLSs into a pseudo-temporal continuum of molecular subtypes—aggregated, activated, and post-activated. Early stages were dominated by niches co-enriched for iCAFs, CCL19⁺ CD8⁺ T cells, and APC-like myeloid subsets, including cDC2s and CXCL12⁺ macrophages, which formed coordinated spatial interaction networks suggestive of TLS initiation and immune recruitment. As TLSs transitioned to the activated state, niches characterized by CXCL13⁺ CD4⁺ T cells and memory B cells emerged as dominant hubs, indicating progression toward germinal center–like immune architecture.

Interestingly, our observation of TLS subtype based on gene expression can only partially align with TLS maturity identified by HE staining morphology. This suggested that traditional clustering strategy for TLS maturity is insufficient to distinguish TLSs which are pre-activated or post-activated. This claimed the necessity to develop more reliable methods for TLS clustering, which was significantly relative to clinical outcome.

To validate and extend these observations, we performed snRNA-seq using the 10x Chromium Flex platform across 14 iCCA samples. This approach enabled fine-grained identification of TLS-associated immune and stromal subsets, including CXCL13⁺ CD4⁺ T cells, memory B cells, and CXCL12⁺ macrophages. Mapping Flex-derived cell types back onto Xenium spatial coordinates using Tangram revealed strong spatial congruence across platforms. Specifically, CXCL12⁺ macrophages and iCAFs were enriched at the periphery of Aggregated TLSs and formed ring-like distributions, while CXCL13⁺ CD4⁺ T cells and memory B cells infiltrated central TLS regions in Activated states. CXCL12⁺ macrophages exhibited high expression of M1-related genes and scored highest for an M1-RTM signature, reinforcing their role as spatially privileged initiators of TLS nucleation. The secretion of CXCL12, TNF, and IL2RA by CXCL12⁺ macrophages may orchestrate immune cell recruitment and spatial organization—CXCL12 guides cDC2 and lymphocyte positioning, TNF potentiates stromal activation and lymphoid chemokine production (e.g., CCL19, we also identified enrichment of CCL19^+^ T cells in Agg subtype), while IL2RA could modulate local T cell survival, collectively reinforcing TLS aggregation. These results support a stepwise model in which chemokine production, antigen presentation and CD4⁺ T cell activation by M1-like RTMs and DCs precede lymphocyte recruitment and follicular organization, ultimately establishing mature TLS architecture—a model consistent with peripheral apc-CN to TLS-score CN transition defined in our IMC analysis.

Several clinical implications emerge from these findings. Our random forest classifier, derived from proteomic features, may serve as a diagnostic tool for TLS-based tumor stratification. More importantly, mechanistic insights into M1-like RTM and apc-CN function suggest opportunities for therapeutic TLS modulation—either by promoting their spatial assembly or by reprogramming immunosuppressive counterparts. In our previous work, we identified M2-like RTMs as mediators of immunotherapy resistance through suppression of CD8⁺ T cells, where their spatial proximity to CD8⁺ cells was associated with poor clinical outcomes in iCCA (*4*). Building on these findings, reprogramming M2-like into M1-like RTMs emerges as a rational strategy to enhance anti-tumor immunity.

This study is not without limitations. Our analyses are primarily observational and descriptive. Future investigations utilizing genetic manipulation models are needed to validate causal relationships between M1-like RTMs and TLS formation. Moreover, longitudinal spatial profiling may clarify whether TLS maturation states represent stable subtypes or transient phases within an evolving tumor-immune ecosystem.

In conclusion, our spatial ecosystem analysis redefines the landscape of TLS biology in iCCA. We show that coordinated cellular reorganization and niche-level remodeling underlie TLS heterogeneity, with implications for both biomarker development and therapeutic targeting. These insights offer a foundation for future strategies aiming to harness or reengineer TLSs to overcome immune resistance in this challenging malignancy.

## Methods

### Histopathological Analysis

All patients in the in-house cohorts were evaluated using standardized protocols. The study was conducted in accordance with the principles of the Helsinki Declaration and received ethical approval from the Medical Ethics Committees of the following institution: Sun Yat-sen University Cancer Center (SYSUCC). This retrospective study analyzed archived formalin-fixed, paraffin-embedded (FFPE) tissue specimens collected for routine clinical purposes. All patient data were anonymized prior to analysis, and the requirement for informed consent was waived by the Ethics Committees. The validation cohort, comprising 155 patients, was described in our previous study (*19*). Hematoxylin and eosin (H&E) staining was performed on 4-μm formalin-fixed, paraffin-embedded (FFPE) iCCA tissue sections following standard protocols. After deparaffinization in xylene (3 × 5 min) and rehydration through graded ethanol series (100%, 95%, 75%, 50%; 5 min each), sections were washed in phosphate-buffered saline (PBS, pH 7.4; Solarbio, cat# P1020). Nuclear staining was achieved using Mayer’s hematoxylin (Solarbio, cat# G1080) for 5-8 min at room temperature, followed by differentiation in 1% acid ethanol and bluing in 1% ammonium hydroxide (Solarbio, cat# G1822). Cytoplasmic counterstaining employed eosin Y (Solarbio, cat# G1100) for 2 min. Processed sections were dehydrated, cleared in xylene, and mounted with resinous medium (Solarbio, cat# G1402). Whole-slide imaging was conducted using a Pannoramic 250 Flash II Scanner (3D HISTECH) at 20× magnification. Two board-certified pathologists, blinded to clinical outcomes, independently reviewed H&E slides to confirm iCCA diagnosis and demarcate tumor regions of interest (ROIs), including tumor core, invasive margin, and immune cell-enriched regions.

### Immunohistochemical Staining

Automated immunohistochemistry (IHC) was performed on 4-μm FFPE sections using the BOND RX system (Leica Biosystems). Following deparaffinization and rehydration, antigen retrieval was performed in EDTA buffer (pH 9.0; Leica, cat# AR9640) at 100°C for 20 min. Endogenous peroxidase activity was blocked with 3% H₂O₂ before incubation with primary antibodies: anti-CD20 (1:200; Abcam ab955), anti-CD21 (1:200; Abcam ab955), anti-CD3 (1:200; Abcam ab955), anti-BCL6 (1:200; Abcam ab955), anti-Ki-67 (1:200; Abcam ab955) for 20 min at 37°C. Detection employed the BOND Polymer Refine Detection Kit (Leica, cat# DS9800) with DAB chromogen development. Counterstaining with hematoxylin (Leica, cat# CS700), dehydration, and mounting preceded whole-slide scanning (Pannoramic 250 Flash II, 20×).

### Sample Preparation and Proteomic Analysis

FFPE tissue samples were processed through sequential dewaxing and protein extraction steps prior to mass spectrometry analysis. Samples were initially equilibrated at 37°C for 30 minutes, followed by two 10-minute heptane incubations for paraffin removal. Gradual rehydration was achieved through a series of ethanol washes (100%, 90%, and 75% ethanol, 5 minutes each). The dewaxed samples were then transferred to PCT tubes containing 12.5 μL of 100 mM Tris-HCl buffer (pH 10) and subjected to protein extraction at 95°C with constant agitation (600 rpm) for 30 minutes.

Protein samples underwent reduction and alkylation in 30 μL of lysis buffer (6 M urea, 2 M thiourea, 100 mM TEAB) supplemented with 5 μL of 200 mM TCEP and 2.5 μL of 800 mM IAA using Pressure Cycling Technology (PCT). This process involved 90 cycles of alternating high (45,000 psi for 30 seconds) and ambient pressure at 30°C. Following buffer dilution with 75 μL of 0.1 M TEAB, enzymatic digestion was performed with trypsin (0.5 μg/μL) and Lys-C (0.25 μg/μL) under 20,000 psi pressure through 120 cycles (50 seconds high pressure/10 seconds ambient pressure) at 30°C. The digestion was terminated by acidification with 15 μL of 10% trifluoroacetic acid, and the resulting peptides were desalted using SOLAμ SPE cartridges (Thermo Fisher Scientific) after confirming sample pH between 2-3.

LC-MS/MS analysis was conducted on a Vanquish Neo UHPLC system coupled to an Orbitrap Astral mass spectrometer (Thermo Scientific). Peptides were separated on a reversed-phase analytical column (1.9 μm, 120 Å pore size, 150 mm length × 75 μm inner diameter) using a 500 nL/min flow rate with a 19-minute gradient from 8% to 40% mobile phase B (0.1% formic acid in 80% acetonitrile). The mass spectrometer operated in data-independent acquisition (DIA) mode with FAIMS voltage set to -42 V. Full scan MS spectra were acquired from 380-980 m/z at 240,000 resolutions, while MS/MS spectra covered 150-2000 m/z with 25% collision energy. The analysis employed 299 isolation windows (2 m/z width) with automatic gain control targets of 500% and maximum injection times of 3 ms for both MS1 and MS2 scans.

Mass spectrometry data were processed using DIA-NN software (version 1.8.1) against the UniProt human reference proteome database (2023-07-25 release) with match-between-runs enabled. Static modification was set for carbamidomethylated cysteine residues, while methionine oxidation was specified as a variable modification. All identifications were filtered at a 1% false discovery rate threshold to ensure data quality.

### Random Forest Model Construction and Validation

We employed the R package ranger to construct a random forest classification model for predicting iTLS status (iTLS^+^ vs. iTLS^-^) based on proteomic profiles. The model was trained on the discovery cohort (n = 214) and subsequently validated on an independent cohort (n = 155).

To ensure robustness, we performed 10 independent training iterations, each initialized with a distinct random seed. In each iteration, the discovery cohort was randomly partitioned into training (75%) and internal testing (25%) subsets. Hyperparameter optimization was conducted via 5-fold cross-validation using a grid search strategy to identify the optimal model configuration. The best-performing parameters from each iteration were used to train the final model, and its predictive performance was evaluated on the held-out test subset using receiver operating characteristic (ROC) analysis. The results from all 10 iterations were aggregated to assess overall model stability and accuracy. The validated model was then applied to the independent validation cohort (n = 155) to evaluate its generalizability. Input data consisted of standardized proteomic expression matrices, with binary iTLS classification as the outcome variable.

### Gene Set Enrichment Analysis and Gene Ontology analysis

Gene Set Enrichment Analysis (GSEA) was performed using the hallmark gene sets from the Molecular Signatures Database (MSigDB) to identify enriched biological pathways across TLS molecular subtypes. The analysis was conducted with GSEA software using 1000 gene set permutations with weighted enrichment statistic and signal-to-noise ratio for gene ranking. Gene sets meeting significance thresholds of false discovery rate (FDR) < 0.25 and nominal p-value < 0.05 were considered statistically enriched. For single-sample GSEA (ssGSEA), we calculated enrichment scores for each sample using the same hallmark gene set collection, which were subsequently correlated with immune cell abundances and clinical parameters. The gene signature used in this study was summarized in supplemental table S1.

Gene Ontology (GO) analysis was performed to identify significantly overrepresented biological processes, molecular functions, and cellular components. We applied Benjamini-Hochberg correction with an adjusted p-value cutoff of 0.05 and required a minimum of 5 differentially expressed genes per term. To reduce redundancy among significant GO terms, semantic similarity analysis was performed. KEGG pathway enrichment analysis was performed using the clusterProfiler R package (*20*), with pathways considered significantly enriched at a false discovery rate (FDR) < 0.05.

### Immune Architecture Analysis

Building on previously established methods (*21*), immune cell interactions were analyzed using a k-nearest neighbor graph built with the *buildSpatialGraph* function from the imcRtools package. The analysis pipeline comprised three key computational steps: First, spatial graphs were constructed using the buildSpatialGraph function. Cellular neighborhoods (CNs) were subsequently identified through k-means clustering (k=20) based on both cellular phenotypes and spatial relationships. Second, cell-cell interactions were quantified using an expanded neighborhood graph (15 μm expansion radius) where physically contacting cells were considered interacting partners. The testInteractions function (histcat method) was implemented to calculate interaction frequencies between all possible cell type pairs. For each cell subset A, we determined the number of adjacent subset B cells and normalized this by the proportion of A cells having ≥1 B-type neighbor within each region of interest (ROI). Statistical significance was assessed through 1,000 permutations of cell type labels while maintaining spatial structure, generating empirical null distributions. Significant interactions (attraction or avoidance) were defined as those exceeding the 95th or falling below the 5th percentile of the null distribution, respectively. Third, minimum intercellular distances were computed using the minDistToCells function to quantify spatial relationships between specific immune subsets. Fourth, effective score was calculated by total counts of a specific neighbor cell type which is aggregated from k-nearest neighbor matrix, and then summarized by mean for each center cell type (*22*). All analyses were performed at the ROI level, with results aggregated by spatial immune type for comprehensive visualization.

Spatial context maps were generated using a custom analysis pipeline developed by Garry P. Nolan’s group (*14*). To assess CN–CN spatial associations, we adapted the general framework used for cellular neighborhood identification with several key modifications. For each cell, spatial windows were defined using the 100 nearest neighbors to capture broader local tissue architecture. Within each window, we used CN labels—rather than single-cell annotations—to quantify the composition.

Instead of clustering the windows, we determined the smallest combination of CNs that collectively accounted for more than 85% of the cells within each window, which we defined as the spatial context. This approach captured the dominant CN interactions in each local area. All unique combinations were counted and aggregated to construct spatial context maps.

In the resulting network graphs, each node represented a unique CN combination, with color indicating the CN pairing and node size reflecting its relative frequency across the tissue. Graph rows corresponded to the number of CNs forming the dominant combination in a window: windows dominated by a single CN (>85%) were placed in the first row; windows with two CNs contributing >85% were placed in the second row; and those with three or more CNs in the third or higher rows.

The number of CNs per spatial context provided insight into underlying structural biology. Single-CN dominance indicated spatial compartmentalization, reflecting large regions with uniform CN composition. Conversely, combinations involving three or more CNs reflected mixed immune environments. Of particular interest, two-CN combinations making up ≥85% of a window suggested structured interfaces between immune compartments that may underlie coordinated functional activity.

### Xenium In Situ Spatial Transcriptomic Analysis

The Xenium In Situ platform (10x Genomics) was employed for high-resolution spatial transcriptomic profiling using a fully integrated system with automated sample handling, liquid processing, and wide-field epifluorescence imaging capabilities. The instrument features a dual-slide configuration with an imaging area of 12 × 24 mm per slide and incorporates an optimized analytical pipeline performing image preprocessing, fluorescent puncta detection, transcript decoding with quality score assignment, and nuclear segmentation based on DAPI staining followed by cytoplasmic expansion to a 15 μm radius or until adjacent cell boundary contact. The platform generates standardized output files including feature-cell matrices in HDF5/MEX formats compatible with 10x Genomics’ single-cell RNA-seq pipelines, transcript localization files containing mRNA coordinates and quality metrics, and CSV-formatted cellular boundary definitions. For region-specific analysis, tissue domains of interest were manually annotated using the polygon annotation tool in Xenium Explorer software (10x Genomics developmental version), with coordinate data exported in CSV format for subsequent computational integration, enabling precise interrogation of transcriptomic patterns within histologically defined tissue compartments.

Niche analysis is based on the BuildNicheAssay function from within Seurat to construct a new assay called niche, which contains the cell type composition spatially neighboring each cell. Due to the multiple fovs in our Xenium data, we first calculate the 25 nearest neighbors in each fov separately and then merge the matrix together for k-means clustering (k = 14).

For network visualization, the minimum distance between each cell type/niche was computed and visualized using the R package qgraph (*23*). Frequency of each cell type/niche was calculated for node size, and cellular/niche neighborhood (k = 25) was aggregated for edge width.

### TLS states identification in Xenium

A total of 61 TLS structures was first identified in the Xenium explorer software regarding on both histological and molecular features. Pseudo-bulk expression matrix for each TLS was generated from AggregateExpression function from Seurat package. Inspired by previously established methods(*24*), we first performed PCA of TLSs in each sample separately. TLSs in 4647T samples shows largest diversity among all samples. Depended on this, we set TLSs from 4647T as a reference and projected TLSs from other samples onto the 4647T’s low-dimension space. Expression level of representative gene including CXCL12, CXCL13, BANK1, CR2, RORC, and IRF8 were characterized. To classify TLSs into different states, we performed k-means clustering on TLSs using the first two principal components (PC1 and PC2) with k set to 3. Together with cell type components, we defined three pseudo-temporal TLS states: Aggregated, Activated, and Post-activated. The cluster with a high LTi signature (RORC) represented the early aggregated state. The cluster with high expression of functional chemokines (CXCL13) and germinal center relative gene (CR2) refers to the activated TLS state. The last cluster with descending chemokine expression and increasing hypoxia signature were considered as post-activated TLS.

### Single-nucleus RNA Sequencing from FFPE Tissue

FFPE tissue curls (50 μm) were dissociated using Liberase™ TH Research Grade (Roche Diagnostics Deutschland GmbH, Cat# 05401151001) following the manufacturer’s protocol. The resulting nuclei were washed, counted, and resuspended, and approximately 16,000 nuclei were loaded per well on a Chromium Chip Q (10x Genomics), targeting a recovery of ∼10,000 nuclei per GEM well.

Library preparation was performed using the Chromium Fixed RNA Profiling Kit (10x Genomics, Cat# 1000474) according to the manufacturer’s instructions. Sequencing was carried out on an Illumina NovaSeq™ X Plus system using paired-end dual indexing. Raw BCL files were demultiplexed using bcl2fastq (Illumina), and the resulting FASTQ files were processed with Cell Ranger v7.0.1 (10x Genomics) using the multi pipeline and aligned to the GRCh38-2020-A human reference genome. Quality control and integration were performed using the “Seurat” R package (version 4.4.0) (*25*). To remove low-quality data, genes expressed in fewer than three cells were excluded. Cells with fewer than 200 or more than 4,000 detected genes, were filtered out to eliminate barcodes corresponding to empty droplets or potential doublets. Cells with mitochondrial gene content exceeding 15% were also excluded. Doublets and multiplets were identified and removed using the scDblFinder algorithm with default parameters (*26*).

For data integration across individuals, Seurat’s *IntegrateData* function was applied to correct for batch effects and project all single cells into a shared low-dimensional space. After integration, a graph-based unsupervised clustering algorithm was implemented using Seurat’s default pipeline. The UMI count matrix was first normalized with the *NormalizeData* function, and log-normalized expression values were used to identify 2,000 highly variable genes via the *FindVariableFeatures* function with the “vst” method. These genes were then used for dimensionality reduction by principal component analysis (PCA), with 20 PCs retained after regressing out total UMI counts.

Clustering was performed on the top 20 PCs using the *FindClusters* function at a resolution of 1.2. Clusters were visualized via Uniform Manifold Approximation and Projection (UMAP) and annotated based on canonical lineage markers. To define cluster-specific marker genes, the *FindAllMarkers* function was applied using the Wilcoxon rank-sum test, with thresholds set to min.pct = 0.25 and logfc.threshold = 0.25. Identified marker genes, together with known lineage-specific signatures, were used for downstream cluster annotation. The 10x flex sequencing data was spatially mapped to 10x xenium data by Tangram package(*27*) at cluster level with default parameters.

### Tissue Distribution of Clusters

For each cell subtype, we evaluated its distribution pattern across the four groups through the calculation of the Ro/e as previously described (*28*). Specifically, Ro/e is the ratio of the observed cell number to the expected cell number of a given combination of cell subtype and tissue. We used the “calTissueDist” function from the Startrac package to obtain the observed and expected cell numbers, using cell frequency as the input data. Briefly, a cell subtype was more frequently observed in a specific tissue than random expectations when Ro/e >1 and was therefore assumed to be enriched in that group.

## Statistical analysis

All statistical analyses were performed using the R software (Version 4.4.1) and the Python software (Version 3.12.7). The Cox proportional-hazards model was utilized to compare the survival outcomes. Data are presented as mean ± standard deviation. Statistical significance was determined as follows: *p < 0.05, **p < 0.01, ***p < 0.001.

## Acknowledgements

This work was supported in part by the National Natural Science Foundation of China 82473471 (to X.B.), 82102817 (to X.D.), 82373428 (to W.F.) and by Zhejiang Provincial National Science Foundation of China LY23H160013 (to X.B.) and by the Major Scientific Project of Zhejiang Province 2023C03061 (to W.F.) and by the “Pioneer” and “Leading Goose” R&D Program of Zhejiang Province 2023C03061 (to W.F.) and 2025C02073 (to P.Z.). We give thanks for technical support by the Central Laboratory (Zhi Jiang Division), the First Affiliated Hospital, School of Medicine, Zhejiang University.

## Author contributions

Liaoliao Gao, Jie Mei, Libing Hong and Yuzhi Jin contributed equally to this work. Xuanwen Bao, Weijia Fang and Wei Wei designed the study. Liaoliao Gao, Jie Mei, Libing Hong, Yuzhi Jin and Jinlin Cheng analyzed and interpretated the data. Jinlin Cheng performed H&E and pathological evaluation. Liaoliao Gao, and Libing Hong performed the visualization work. Liaoliao Gao and Xuanwen Bao wrote the manuscript. Xuqi Sun, Xiaomeng Dai, Bo Lin, Yajie Sun, Peng Zhao, Rongping Guo, Jingping Yun, Minshan Chen and Inmaculada Martínez-Reyes edited the manuscript. All the authors read and approved the final manuscript.

## Data availability

The proteomics data have been deposited in the iProX under the project number IPX0011444000. The dataset will be made publicly available upon manuscript acceptance (https://www.iprox.cn/page/PSV023.html;?url=17440935604958xLu; password: hVtW). The snRNA-seq data have been deposited in the GDSC under the project number HRA012124. The dataset will be made publicly available upon manuscript acceptance. Publicly available datasets used in this study can be accessed from the Gene Expression Omnibus (GEO) under accession numbers GSE91061 and GSE126044, and from the European Nucleotide Archive (ENA) under accession number PRJEB23709.

