## Supplementary Figs. S1 to S7 and Supplementary Tables S1 to S3. for "Spatiotemporal Mapping of Tertiary Lymphoid Structure Heterogeneity Shapes Immune Niches and Clinical Outcomes in Intrahepatic Cholangiocarcinoma"

#### Supplementary Text

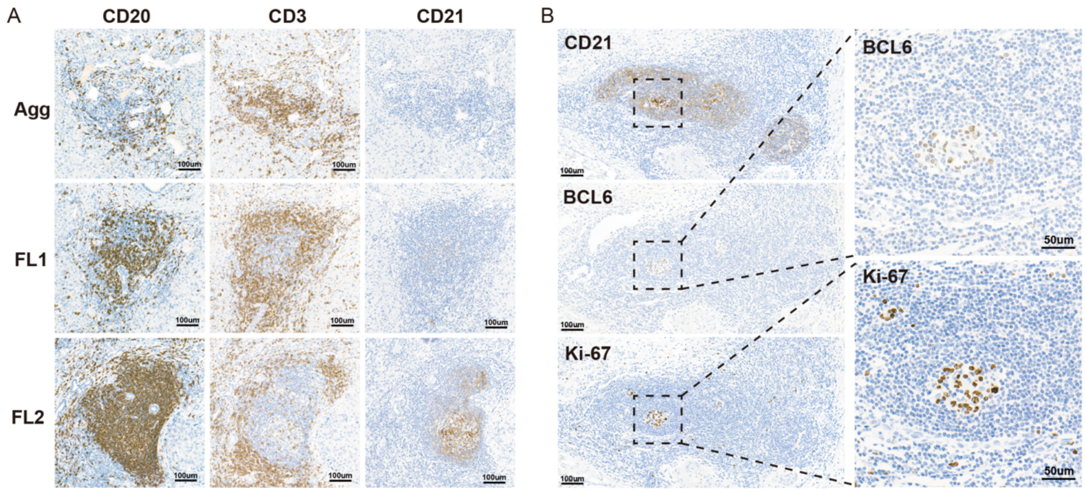


**Fig. S1. Histological characterization of TLS morphologies.**

(A) Representative IHC staining of CD20, CD3, and CD21 in TLS subtypes: aggregates (AGG), primary follicles (FL1), and secondary follicles (FL2). Scale bar = 100 µm.

(B) A typical TLS, with IHC staining showing CD21, BCL6, and Ki67 expression. The magnified core region highlights the BCL6 and Ki67 staining. Scale bar = 100 µm and 50 µm, respectively.

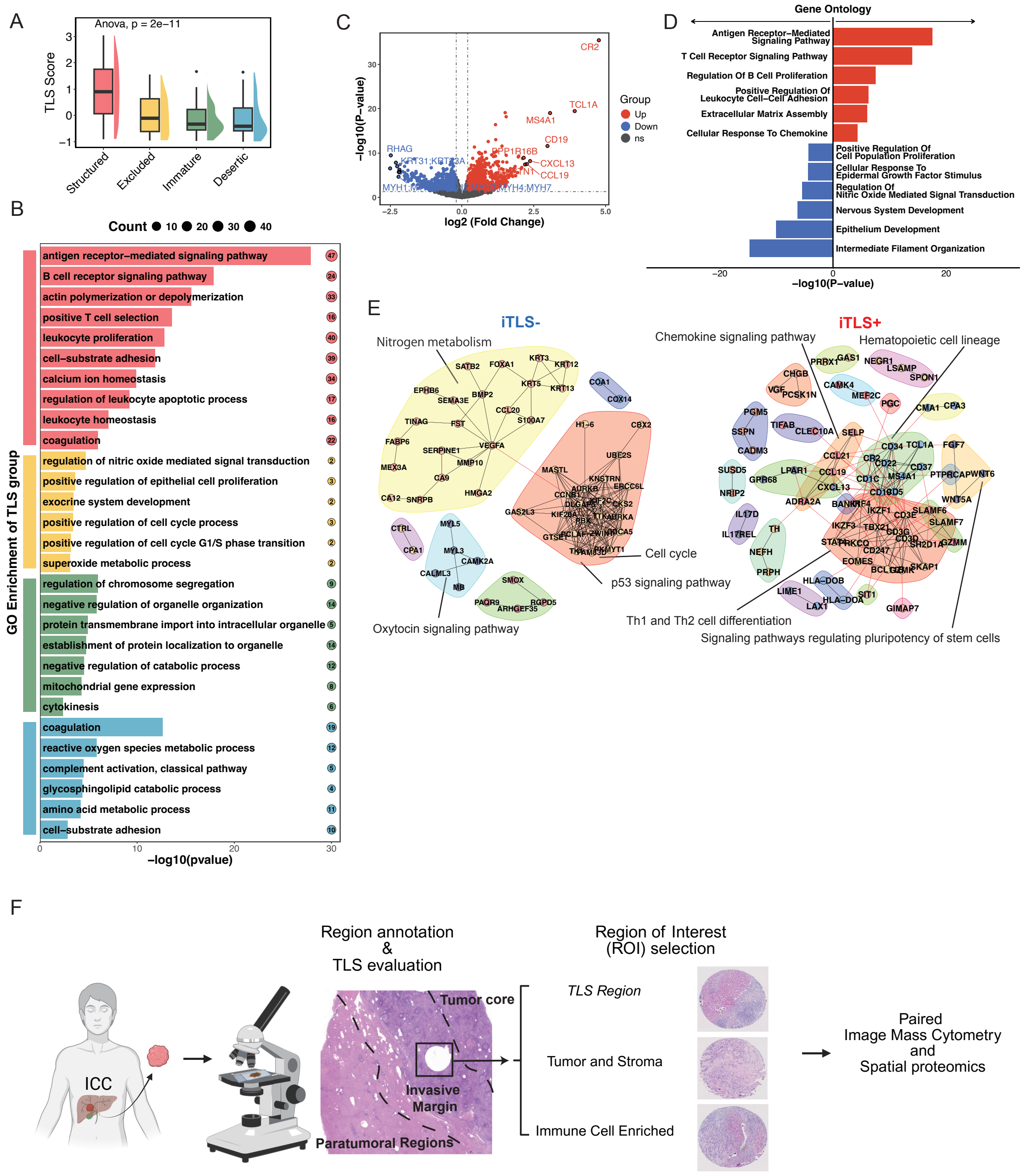
A
Anova, p = 2e−11
TLS Score
Structured
Excluded
Immature
Desertic
B
Count ● 10 ● 20 ● 30 ● 40
GO Enrichment of TLS group
antigen receptor−mediated signaling pathway
B cell receptor signaling pathway
actin polymerization or depolymerization
positive T cell selection
leukocyte proliferation
cell−substrate adhesion
calcium ion homeostasis
regulation of leukocyte apoptotic process
leukocyte homeostasis
coagulation
regulation of nitric oxide mediated signal transduction
positive regulation of epithelial cell proliferation
exocrine system development
positive regulation of cell cycle process
positive regulation of cell cycle G1/S phase transition
superoxide metabolic process
regulation of chromosome segregation
negative regulation of organelle organization
protein transmembrane import into intracellular organelle
establishment of protein localization to organelle
negative regulation of catabolic process
mitochondrial gene expression
cytokinesis
coagulation
reactive oxygen species metabolic process
complement activation, classical pathway
glycosphingolipid catabolic process
amino acid metabolic process
cell−substrate adhesion
−log10(pvalue)
C
−log10(P−value)
log2 (Fold Change)
Group
Up
Down
ns
CR2
TCL1A
MS4A1
CD19
CXCL13
CCL19
RHAG
D
Gene Ontology
Antigen Receptor−Mediated Signaling Pathway
T Cell Receptor Signaling Pathway
Regulation Of B Cell Proliferation
Positive Regulation Of Leukocyte Cell−Cell Adhesion
Extracellular Matrix Assembly
Cellular Response To Chemokine
Positive Regulation Of Cell Population Proliferation
Cellular Response To Epidermal Growth Factor Stimulus
Regulation Of Nitric Oxide Mediated Signal Transduction
Nervous System Development
Epithelium Development
Intermediate Filament Organization
−log10(P−value)
E
iTLS-
iTLS+
Nitrogen metabolism
Chemokine signaling pathway
Hematopoietic cell lineage
Cell cycle
p53 signaling pathway
Oxytocin signaling pathway
Th1 and Th2 cell differentiation
Signaling pathways regulating pluripotency of stem cells
F
Region annotation
&
TLS evaluation
Region of Interest
(ROI) selection
ICC
Tumor core
Invasive
Margin
Paratumoral Regions
TLS Region
Tumor and Stroma
Immune Cell Enriched
Paired
Image Mass Cytometry
and
Spatial proteomics

**Fig. S2. Proteomic validation of TME subtypes and iTLS-associated molecular programs.**

(A) TLS score across four proteomics-defined immune subtypes (Structured, Immature, Excluded, Desertic), calculated using a previously published TLS gene signature applied to bulk proteomic data. Structured tumors showed the highest TLS scores, while Desertic tumors had the lowest.

(B) Gene Ontology (GO) enrichment analysis of differentially expressed proteins across TME subtypes. Structured tumors showed upregulation of immune-related pathways such as B cell receptor signaling, leukocyte proliferation, and T cell selection. Desertic tumors were enriched for coagulation, ROS metabolic processes, complement activation, and ECM remodeling (e.g., cell–substrate adhesion).

(C) Volcano plot comparing protein expression between $iTLS^+$ and $iTLS^-$ tumors in the discovery cohort. Key TLS-associated markers, including CR2, MS4A1, CD19, CXCL13, and CCL19, were significantly upregulated in $iTLS^+$ tumors.

(D) GO enrichment analysis of differentially expressed proteins between $iTLS^+$ and $iTLS^-$ tumors. $iTLS^+$ samples showed enrichment in antigen receptor–mediated signaling, T cell receptor pathways, and B cell proliferation, while $iTLS^-$ tumors were enriched for ECM reorganization and metabolic processes.

(E) KEGG pathway analysis revealed $iTLS^+$ tumors were enriched for immune-related pathways such as chemokine signaling and Th1/Th2 cell differentiation, whereas $iTLS^-$ tumors showed enrichment of p53 signaling, cell cycle, and metabolic stress pathways.

(F) Graphical summary of definition and consistency of ROIs for spatial proteomics.

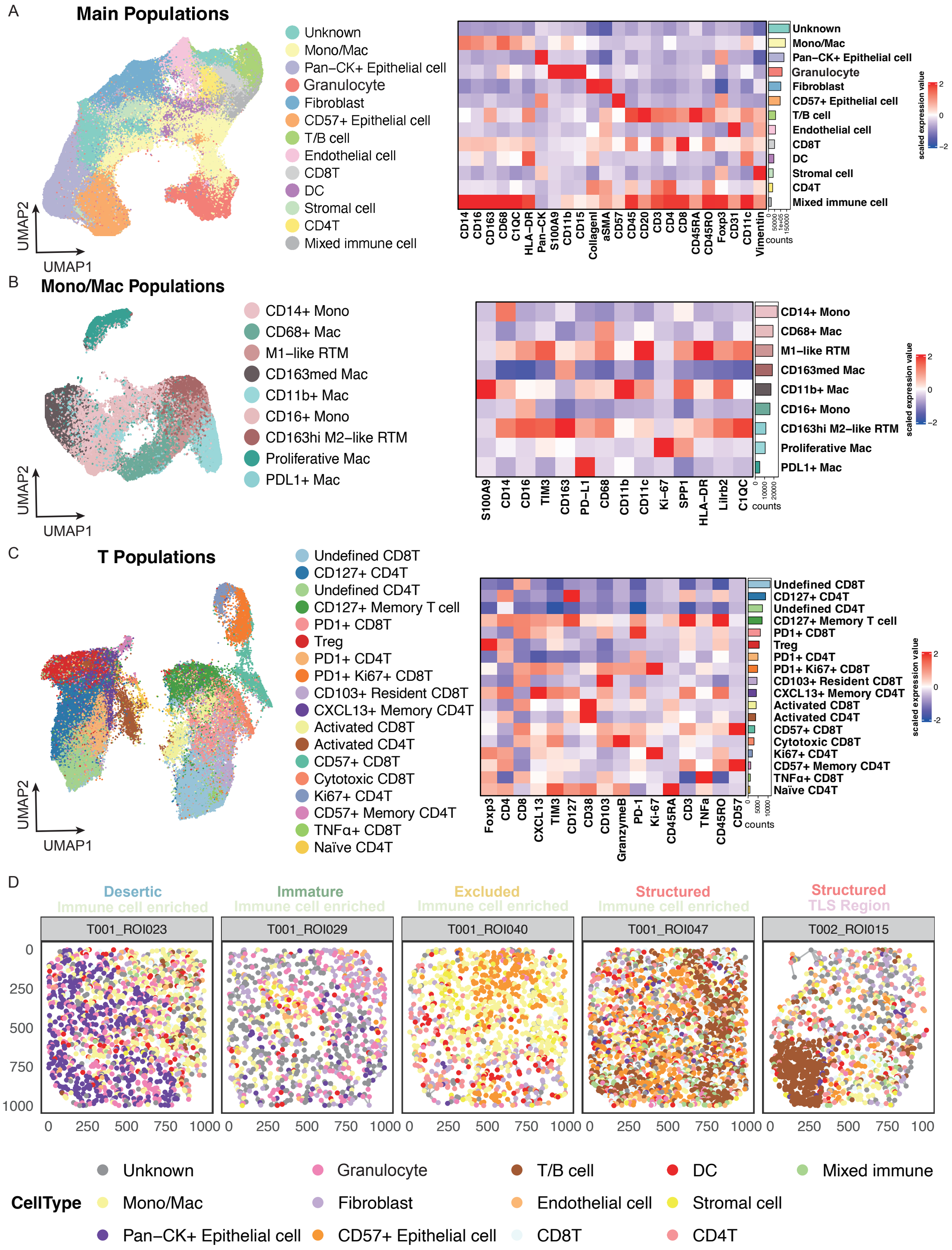


**Fig. S3. Single-cell clustering and immune marker characterization across TME subtypes.** (A) UMAP showing major cell population classification with corresponding IMC marker expression.

(B) UMAP of monocyte/macrophage subsets and corresponding IMC marker expression.

(C) UMAP of T cell subsets and corresponding IMC marker expression.

(D) Representative ROE-selected immune-enriched ROIs from four TME subtypes, alongside a structured TLS region, highlighting typical cellular compositions.

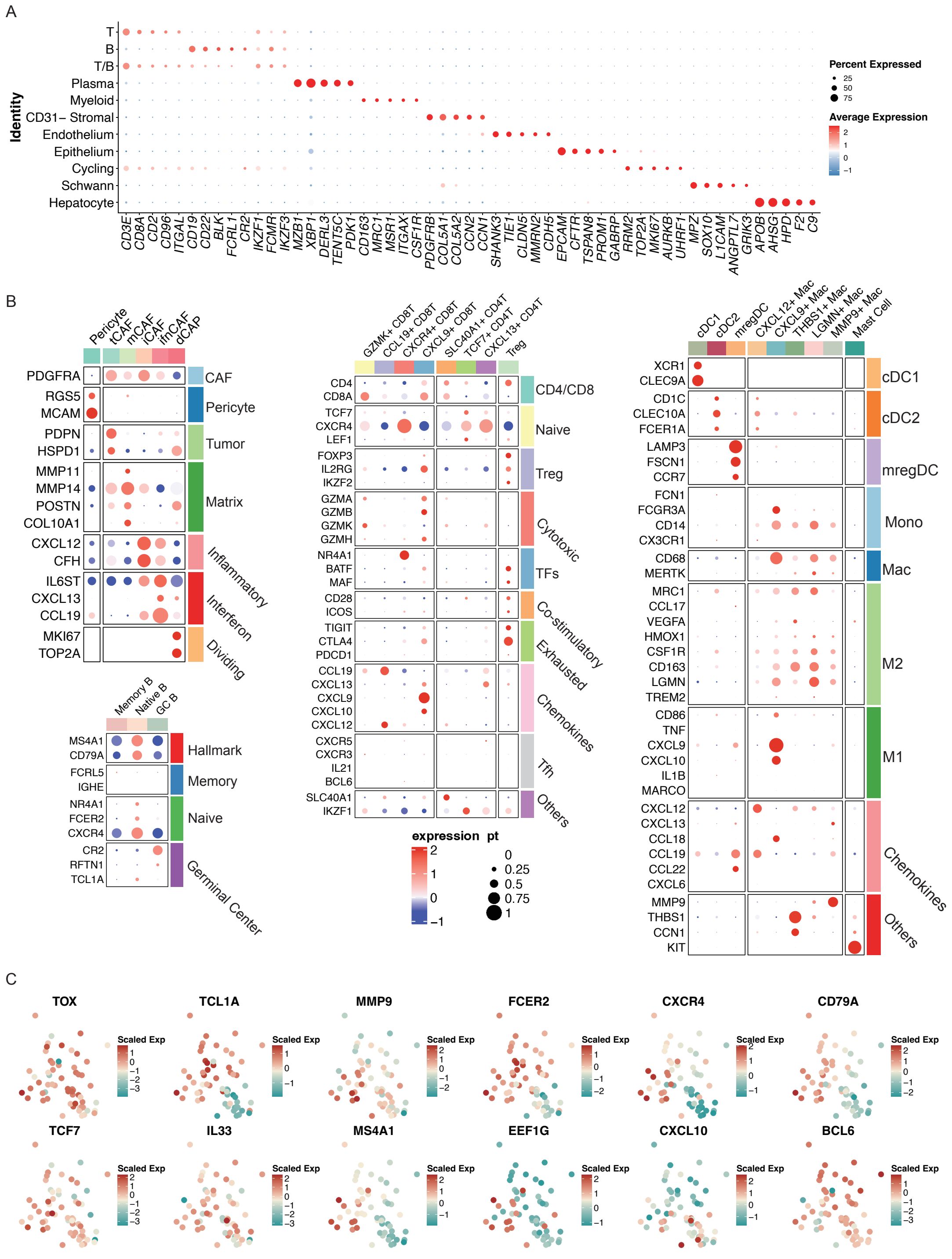


**Fig. S4. Marker gene validation and TLS transcriptional heterogeneity.**

(A) Dot plot showing the top five marker genes for each major cell lineage cluster (e.g., T cells, B cells, myeloid cells, stromal cells) in the Xenium spatial transcriptomic dataset.

(B) Subtype-specific gene signatures across immune and stromal compartments, including hallmark, naïve, memory, and germinal center markers for B cells; lineage, naïve, cytotoxic, Tfh, exhausted, and chemokine-related genes for T cells; and lineage-specific (DC/monocyte/macrophage), M1/M2 polarization, and chemokine genes for myeloid cells.

(C) PCA plot displaying single-TLS expression of canonical TLS-associated genes (e.g., CXCL13, CR2, BANK1), extending analysis in Figure 5F to highlight transcriptional diversity across individual TLSs.

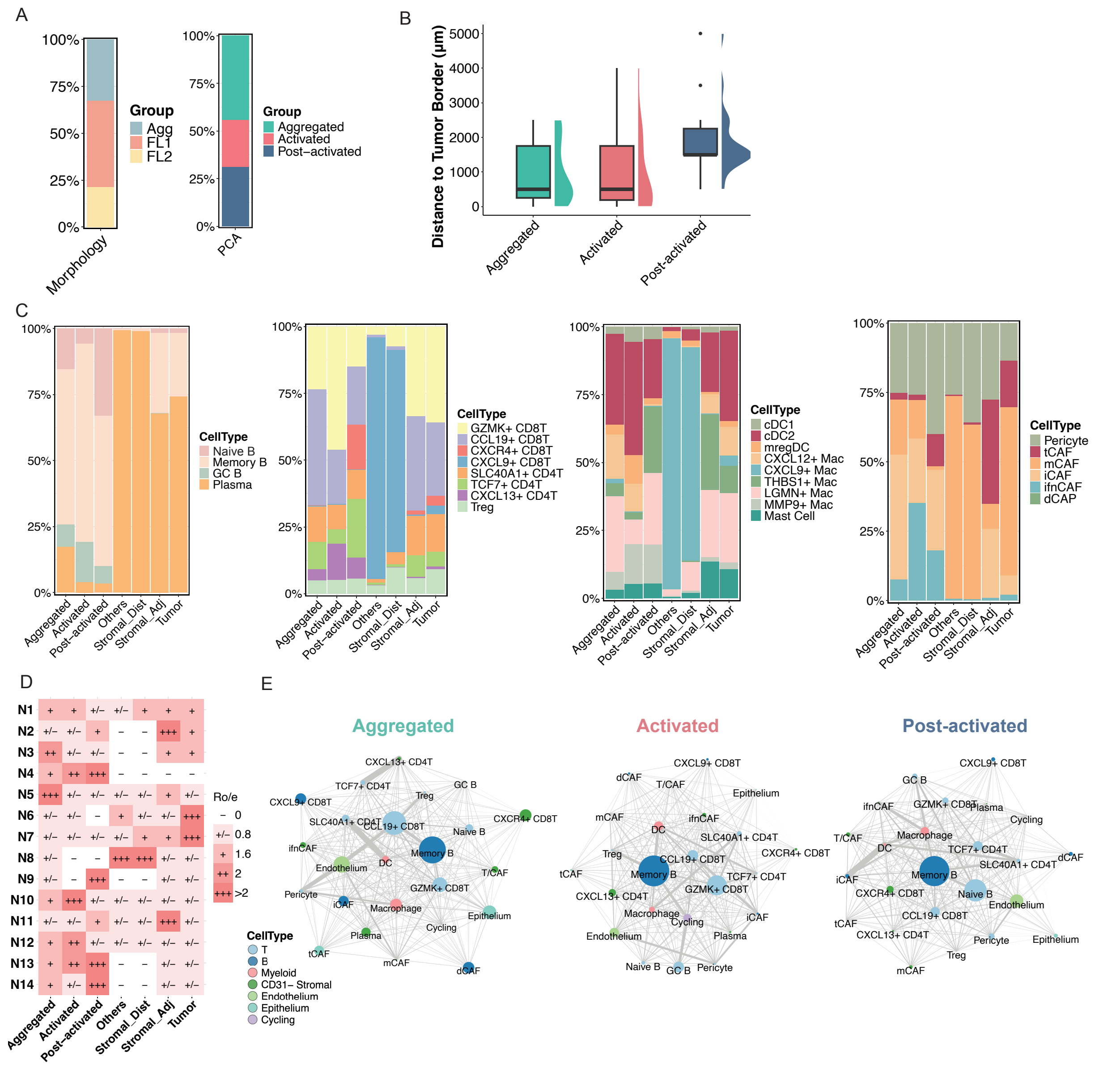
A
100%
75%
50%
25%
0%
Morphology
Group
Agg
FL1
FL2
PCA
Group
Aggregated
Activated
Post−activated
B
Distance to Tumor Border (μm)
5000
4000
3000
2000
1000
0
Aggregated
Activated
Post-activated
C
CellType
Naive B
Memory B
GC B
Plasma
CellType
GZMK+ CD8T
CCL19+ CD8T
CXCR4+ CD8T
CXCL9+ CD8T
SLC40A1+ CD4T
TCF7+ CD4T
CXCL13+ CD4T
Treg
CellType
cDC1
cDC2
mregDC
CXCL12+ Mac
CXCL9+ Mac
THBS1+ Mac
LGMN+ Mac
MMP9+ Mac
Mast Cell
CellType
Pericyte
tCAF
mCAF
iCAF
ifnCAF
dCAP
Aggregated
Activated
Post-activated
Others
Stromal_Dist
Stromal_Adj
Tumor
D
N1 + + +/− +/− + + +
N2 +/− +/− + − − +++ +
N3 ++ +/− +/− − − + +
N4 + ++ +++ − − − −
N5 +++ +/− +/− +/− +/− + +/−
N6 +/− +/− − + +/− +/− +++
N7 +/− +/− +/− +/− + + +++
N8 +/− − − +++ +++ +/− +/−
N9 +/− − +++ − − +/− +/−
N10 + +++ +/− +/− +/− +/− +/−
N11 +/− +/− + +/− +/− +++ +/−
N12 + ++ +/− +/− +/− +/− +/−
N13 + ++ +++ − − +/− +/−
N14 + +/− +++ − − +/− +/−
Ro/e
− 0
+/− 0.8
+ 1.6
++ 2
+++ >2
Aggregated
Activated
Post-activated
Others
Stromal_Dist
Stromal_Adj
Tumor
E
Aggregated
Activated
Post-activated
CellType
T
B
Myeloid
CD31− Stromal
Endothelium
Epithelium
Cycling

**Fig. S5. Spatial distributions and cell–cell interaction networks across TLS molecular subtypes.**

(A) Distribution of TLS morphological and transcriptional maturation states in single-cell spatial transcriptomics.

(B) Spatial proximity of TLS maturation states to tumor boundaries.

(C) Bar plots showing the relative abundance of major cell lineages (B cells, T cells, myeloid cells, CAFs) across TLS subtypes (Aggregated, Activated, Post-activated) and adjacent tissue regions (stromal_distant, stromal_adjacent, tumor).

(D) RO/e analysis showing the tissue privilege of niches across TLS subtypes (Aggregated, Activated, Post-activated) and adjacent tissue regions (stromal_distant, stromal_adjacent, tumor).

(E) Cell–cell interaction networks reconstructed from ligand–receptor analysis for each TLS subtype. Each node represents a cell population; edge thickness corresponds to inferred signaling strength. Cell-level networks recapitulate niche-level rewiring observed in Figure 6C and illustrate subtype-specific communication topologies across TLS maturation stages.

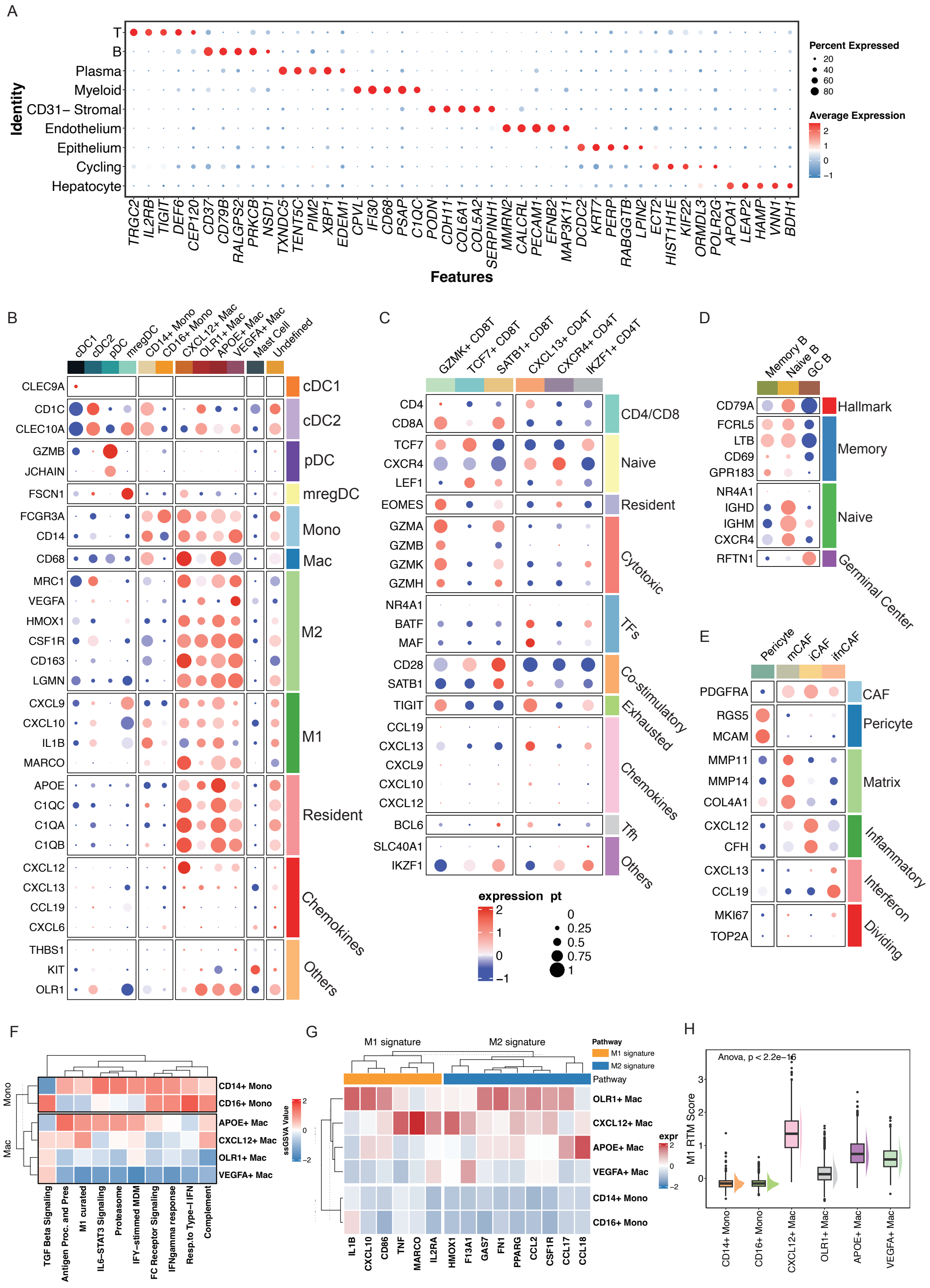


### Fig. S6. Cross-Platform Validation and Subset Characterization of snRNA-seq

(A) Dot plot showing top 5 marker genes for major cell clusters identified by 10x Flex.

(B) Myeloid subset-defining genes, including DC, monocyte/macrophage, M1/M2, residency, and chemokine markers.

(C) T cell signatures: lineage, naive, residency, cytotoxic, exhaustion, TFH, and co-stimulatory genes.

(D) B cell programs: Hallmark, memory, naive, and germinal center markers.

(E) CAF-associated genes.

(F) Heatmap of pathway activity in monocytes and macrophages.

(G) M1 and M2 gene expression across macrophage subsets.

(H) Box plot comparing M1-RTM signature scores, confirming $CXCL12^{+}$ macrophages as the dominant M1-like RTM population.

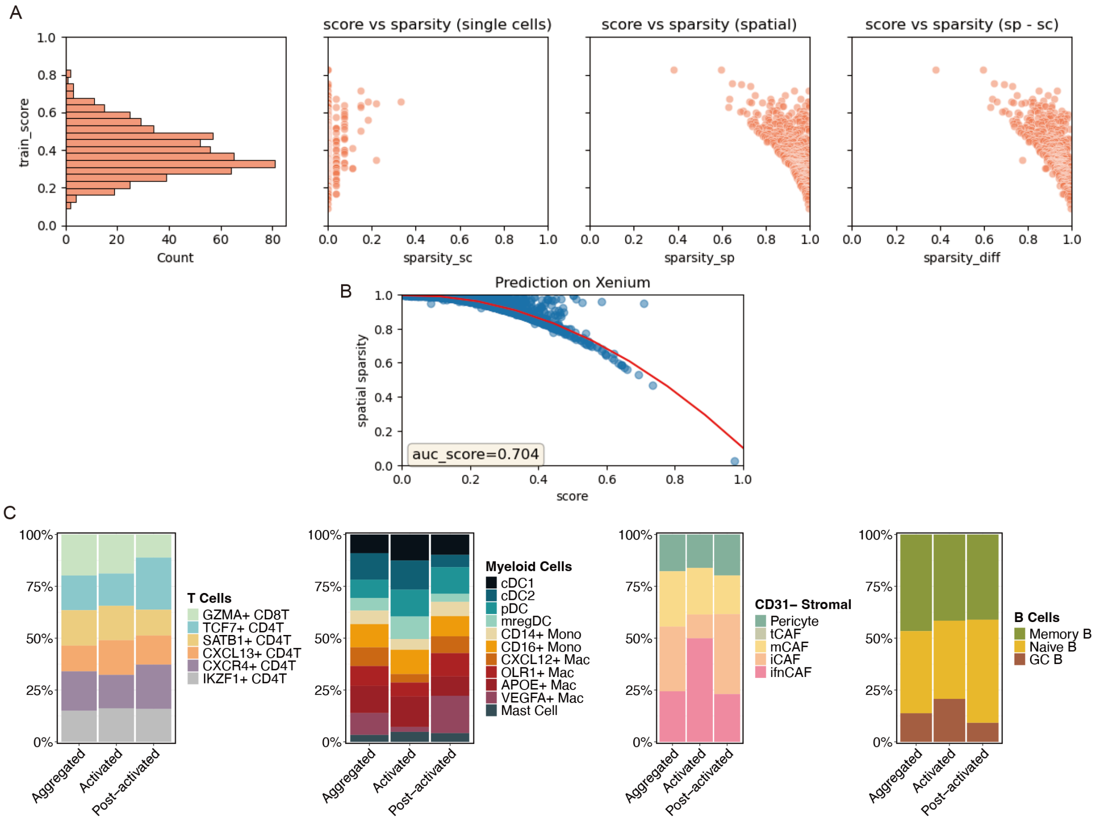


**Fig. S7. Quality control and cell-type mapping validation of Tangram projection.**

(A) Evaluation of Tangram model performance, showing distribution of training scores across cells (left), and scatterplots of matching score versus sparsity in single cells (sparsity_sc), spatial pixels (sparsity_sp), and their difference (sp - sc). These metrics evaluate mapping confidence and exclude overfitting or ambiguous projections.

(B) Spatial prediction accuracy plotted against sparsity across Xenium pixels. Area under the curve (AUC = 0.704) reflects the global performance of the Tangram projection.

(C) Cell-type compositions in Aggregated, Activated, and Post-activated TLS subtypes based on mapped snRNA-seq data. Proportions of T cell, myeloid, stromal, and B cell subsets show expected biological patterns, confirming mapping fidelity.

#### Supplementary Tables

**Table S1: Total signature gene set.**

| Signature | Gene list |
|---|---|
| TLS-related gene signature (REF) | CCL19, CCL21, CXCL13, CCR7, CXCR5, SELL, LAMP3 |
| M1-RTM | C1QC, C1QB, C1QA, AKR1B1, SLC40A1, CCL18, SELENOP, HLA-DOA, RGS1, Z98257.1, SLCO2B1, IGSF6, RHOB, RASSF4, CMKLR1, CHRNA1, SLAMF7, WWC1, ADORA3, AXL, SUCNR1, SDC3, SLCO2A1, SOX18, COL5A3, PXDC1, ADAP2, TMEM37, C2, APLNR, PTMS, NTM, GFPT2, FUCA1, NUAK1, FGF7, ALDH1A1, CNTNAP2, ABCA9, TLR7, SLAMF8, B3GNT7, HES1, SIGLEC8, C1orf54, AQP5, CYP26B1, FABP3, STMN1, CEBPA, OLFML2A, OLFML3, HOXA7, METTL7A, KLF2, CLTRN, SERPINF1, RBP1, EGR1, ESAM, MLPH, MPEG1, RNASE6, MMP10, SMOC1, FAM227A, FUT1, GPR34, BACE2, NEURL3, NUF2, EPS8L1, SEMA3A, SLC7A8, CKS2, CRHBP, IGHM, CFD, ID3, CLIC2, CHI3L2, FOLR2, APOL1, IRF6, CAPN8, PRSS22, ANKRD37, AADAC, TNFSF9, SLC1A3, DST, KIF23, SERPING1, SGO2, CLDN3, EBI3, ANXA13, SEMA6A, OSMR, ENTPD1, EPHX1, CCDC3, MYCL, DLL4, BATF, IL2RA, P2RY13, SCD5, SLC4A1, VSIG2, ASRGL1, CLIC3, MEST, OCLN, TIMM8A, IGHG1, CENPA, GPRC5B, PEG10, HEYL, SYBU, KRTCAP3, LILRB5, TCHH, HGF, RAPGEF5, SLC39A14, CHRM3, LGI4, CCDC68, TMEM119, S100A1, POU2AF1, EBF1 |
| N3-marker | CXCL121, CFH1, DPT, ENG, TIMP21, SLC40A1, COL5A12, AEBP11, MMP142, PDGFRA1, ALDH21, IL1R11, CD3021, FCGR2A1, TNXB, APOL11, MRC22, NRP11, ASPN, PROS1, ADGRA2, EPHA3, INMT, THBS21, AXL2, ADAMTS21, HGF, MSR11, CFHR1, DKK3, CD141, CLEC10A, LGMN1, ITGB51, MYH10, NGFR, DEPTOR, SLC29A1, CFB1 |

| | |
|---|---|
| N4-marker | MS4A1, CR2, CD79A, CXCL131, CD19, TNFRSF13C, CD22, NCF1, CD79B, BANK1, BCL11A, PTK2B, FCRL1, POU2AF1, BLK, FCER2, SERPINA31, STX7, TCL1A, IRF8, BTK, PAX5, PARP1, BTLA, IL4R, RHOH, RUBCNL, CD83, CD200, SNX29, PLEKHA2, CD72, PLCG2, TFEB, RFTN1, WDFY4, CR1, BACH2, CNR2, IFT57, ORAI2, ITIH5, FCRL2, VCAM1, CD40, STAP1, SPIB, CDCA7L, CXCR5, ABCB4, S1PR3, BLNK, TLR10, NT5E |
| N5-marker | CXCL122, DPT1, SLC40A11, CFH2, ENG1, AEBP12, PDGFRA2, TNXB1, INMT1, FBLN52, XBP12, SLAMF71, LGMN2, PROS11, NGFR1 |
| N8-marker | CXCL10, CXCL9, GBP1, WARS, NQO1, CXCL11, IFIT3, IDO1, CLDN18, DHRS2, STAT1, LAD1, UBD, DDR1, TOP2A, EPCAM, SOX9, SKA3, MKI67, CLDN7, GBP5, EPS8L3, PHGDH, TPX2, ST14, HNF4A, CCNE1, CDC20, UBE2T, TAP1, CDK1, EHF, BIRC5, TRPM2, LRRC59, FAT1, TYMP, TRIB3, OAS3, CXCL16, SDHB, POTEF, TONSL, IRF1, MCM4, CSNK2B, ICAM1, KPNA2, GOT1, GRN, AURKB, LAPTM4B, CCT7, MYBL2, APOL6, LAMP3, EIF4A3, CD300A, UGDH, CPVL, PROM1, CSE1L, PRRC2A, ZC3HAV1L, NPM3, NR1H3, ATP2A2, SORD, POSTN, ECHS1, TNFRSF11A, GALE, RCC1, IFIH1, PDIA4, TOX, MTOR, TYMS, CD68, CTSC, CDR2L, LAG3, CCT3, GSTO1, MAD2L1, AIFM1, PRR11, PER3, EZH2, OAS1, NME1, PAPSS1, CFB, SLAMF7, MOV10, PAK1, AHCY, CBWD2, TUBB, FGR, GMDS, TNFSF10, MTCH2, PELO, MAOA, FOXP4, CDK9, EIF2AK2, H2AFX, ANO1, PIK3CB, MX1, STMN1 |
| N10-marker | SOX2-OT1, CCL192, CXCL92, CD8A3, GZMK, MMP9, GZMA, GBP13, CST7 |
| N12-marker | SOX2-OT, CD8A, CD3E, CCL19, GZMK, CD2, SOST, CXCL12, CD96, GIMAP4, SLAMF7, HOXB7, DPT, AOAH, SH2D1A, TC2N, CDC27, FYN, NUCB2, OLFML3 |

| | |
|---|---|
| N13-marker | MS4A11, CD79A1, CD191, TNFRSF13C2, FCMR1, BANK12, CD221, BLK1, CD79B1, BCL11A1, SCIMP1, PARP151, FCRL12, FCRL21, RUBCNL2, PAX51, KCNN4, WDFY41, BTLA1, TFEB1, CNR21, FCRL31 |
| N14-marker | CD3E, IKZF1, TCF7, GIMAP4, LEF1, ITK, CD247, TXK, GIMAP5, CTLA4, ZAP70, CD5, TNFSF8, CD28, PIM1, JAK3, CD6, CD3G, SELPLG, SOCS1, RAPGEF6, SELL, PRDM1, PRKCH, CCND3, SORL1, LEPROTL1 |

**Table S2: Clinical characteristics of the discovery cohort (n=214) for bulk proteomics and H&E staining.**

| | Count | Percentage |
|---|---|---|
| **Gender** | | |
| Female | 84 | 39.25% |
| Male | 130 | 60.75% |
| **Age** | | |
| < = 60 | 99 | 46.26% |
| > 60 | 115 | 53.74% |
| **TNM Stage** | | |
| I | 101 | 47.20% |
| II | 44 | 20.56% |
| III | 60 | 28.04% |
| IV | 9 | 4.21% |
| **Intrahepatic Metastasis** | | |
| 0 | 153 | 71.50% |
| 1 | 61 | 28.50% |
| **Regional Lymph Node Metastasis** | | |
| 0 | 153 | 71.50% |
| 1 | 61 | 28.50% |
| **Perineural Invasion** | | |
| 0 | 164 | 76.64% |
| 1 | 50 | 23.36% |
| **Vascular Invasion** | | |
| 0 | 139 | 64.95% |
| 1 | 71 | 33.18% |
| NA | 4 | 1.87% |
| **Distal Metastasis** | | |
| 0 | 205 | 95.79% |
| 1 | 9 | 4.21% |
| **HBV Status** | | |
| 0 | 121 | 56.54% |
| 1 | 93 | 43.46% |
| **AFP (ng/mL)** | | |
| < = 10 | 176 | 82.24% |
| > 20 | 34 | 15.89% |
| NA | 4 | 1.87% |
| **CA199 (U/mL)** | | |
| < = 37 | 104 | 48.60% |
| > 37 | 101 | 47.20% |

| | | |
|---|---|---|
| NA | 9 | 4.21% |

**Table S3: Clinical characteristics of the validation cohort (n=155) for IMC and spatial proteomics.**

| | Count | Percentage |
|---|---|---|
| **Gender** | | |
| Female | 94 | 60.65% |
| Male | 61 | 39.35% |
| **Age** | | |
| < = 60 | 84 | 54.19% |
| > 60 | 71 | 45.81% |
| **TNM Stage** | | |
| I | 87 | 56.13% |
| II | 25 | 16.13% |
| III | 35 | 22.58% |
| IV | 8 | 5.16% |
| **Differentiation** | | |
| Well | 1 | 0.65% |
| Well to Moderately | 2 | 1.29% |
| Moderately | 45 | 29.03% |
| Moderately to Poorly | 76 | 49.03% |
| Moderately or Poorly | 2 | 1.29% |
| Poorly | 21 | 13.55% |
| NA | 8 | 5.16% |
| **Intrahepatic Metastasis** | | |
| 0 | 117 | 75.48% |
| 1 | 38 | 24.52% |
| **Regional Lymph Node Metastasis** | | |
| 0 | 117 | 75.48% |
| 1 | 38 | 24.52% |
| **Perineural Invasion** | | |
| 0 | 124 | 80.00% |
| 1 | 31 | 20.00% |
| **Vascular Invasion** | | |
| 0 | 106 | 68.39% |
| 1 | 49 | 31.61% |
| **Distal Metastasis** | | |
| 0 | 147 | 94.84% |
| 1 | 8 | 5.16% |
| **HBV Status** | | |
| Negative | 82 | 52.90% |
| Positive | 73 | 47.10% |
| **Tumor Size Diameter (cm)** | | |

| | | |
|---|---|---|
| < = 5 | 84 | 54.19% |
| > 5 | 71 | 45.81% |
| **AFP (ng/mL)** | | |
| < = 10 | 130 | 83.87% |
| > 10 | 25 | 16.13% |
| **CA199 (U/mL)** | | |
| < = 37 | 89 | 57.42% |
| > 37 | 66 | 42.58% |
| **Survival Outcome** | | |
| Deceased in 1 year | 22 | 14.19% |
| Deceased between 1 and 2 years | 26 | 16.77% |
| **Recurrence** | | |
| Recurrence in 1 year | 65 | 41.94% |
| Recurrence between 1 and 2years | 22 | 14.19% |
| No Recurrence | 65 | 41.94% |
| Unknown | 3 | 1.94% |
